# Cell adhesion molecules play subclass-specific roles in electrophysiological response and Schizophrenia risk

**DOI:** 10.1101/2022.11.11.516161

**Authors:** Xinzhe Li, Hongru Hu, Nelson Johansen, Gerald Quon

## Abstract

Multimodal assays such as Patch-seq that simultaneously profile molecular and cellular phenotypes of cells enable the identification of molecular underpinnings of electrophysiological response patterns in neurons. Here we analyzed Patch-seq measurements of thousands of mouse interneurons to identify subclass-specific genes associated with different electrophysiological features. We found extensive subclass specificity: even for the same ephys feature, largely unique sets of genes are associated with that feature in different subclasses. Well established ephys genes such as Reln demonstrated subclass specificity that was previously not reported. Surprisingly, we found that ion channels explained significantly less variation in ephys response across interneurons compared to other genes; in particular, gene sets enriched in cell adhesion genes were amongst the most associated. We found our gene sets associated with action potential dV/dt measurements explained significant heritability of Schizophrenia risk, suggesting a novel role of single neuron electrophysiology in Schizophrenia risk. Finally, we observed significant ephys function switching of cell adhesion molecules across subclasses; the same adhesion molecule was observed to associate with different functional ephys measurements in distinct subclasses and co-express with different genes, suggesting re-purposing of adhesion molecules in different subclasses. Overall, our results yield novel insight into the specificity of roles that individual genes and adhesion molecules play in both single neuron ephys response and Schizophrenia risk.

## INTRODUCTION

Genome-wide association studies (GWAS) have collectively identified hundreds of genetic loci associated with psychiatric disorders, including schizophrenia (SCZ)^1–3^, bipolar disorder (BP)^3–5^, major depressive disorder (MDD)^6–8^, autism spectrum disorder (ASD)^9^, attention deficit hyperactivity disorder (ADHD)^10^, anorexia nervosa^11^, obsessive-compulsive disorder (OCD)^12^, posttraumatic stress disorder (PTSD)^13^ and Tourette’s syndrome (TS)^14^. A central challenge is to characterize the effects of these psychiatric variants on the molecular, cellular and physiological state of cells. At the molecular level, psychiatric variants are enriched in non-coding regions of the genome^15^ such as neuronal regulatory elements^16,17^, and some psychiatric genes have established roles in neuronal plasticity and post-synaptic signalling^18^. At the individual variant level, -omics analyses have identified subsets of psychiatric variants likely to drive changes in gene expression levels and splicing of transcripts that spans across both developmental stages and adulthood^19–21^. On-going efforts are identifying other mechanisms by which variants impact gene regulation, such as through detection of chromatin QTLs, cell fraction QTLs, and chromatin conformational changes^22^.

Psychiatric disorders have also been associated with changes in diverse cellular phenotypes such as cell adherence^23–25^, cellular energetics^26,27^, and neuron electrophysiological response patterns^26,28^. In a limited number of cases, causal links between psychiatric genetic variants and changes in cellular phenotypic variation have been established. With respect to neuron electrophysiology, deletions in the gene SHANK3 leads to an increase of neuronal input resistance and excitability^29^, and 4-bp frameshift mutations in the gene DISC1 yields deficits in presynaptic release. Both genes are linked directly to SCZ^30–33^. However, for the remaining hundreds of genes implicated in psychiatric disorders, it is unclear what roles, if any, those genes may play in driving variation in cellular phenotypes such as single neuron electrophysiology. There are a number of challenges that prohibit genome-wide characterization of genetic variants that are associated with changes in cellular phenotypes. First, unlike most molecular assays that can use post-mortem brain samples^19,20,34–38^, cellular assays such as patch clamping require live neurons that are difficult to obtain from humans *in vivo*. As a result, human studies of neuron function in psychiatric disorders are largely limited to *in vitro* iPSC-derived neurons^39,40^, which are notoriously challenging to efficiently and reproducibly derive without extensive genomic instability^41–48^. Second, the labor-intensive nature of assays such as patch clamping^49^ have prohibited cellular phenotyping of case-control cohort-scale numbers of iPSC-derived neurons, and therefore has inhibited our ability to identify genetic variation associated with cellular phenotypic variation.

To probe the relationship between genetic variation, transcriptomics, and electrophysiology, we performed an unbiased, genome-wide search for genes associated with single neuron electrophysiological (ephys) response to stimuli and used these genes to quantify the heritability of psychiatric disorders with respect to electrophysiological response. Across 52 ephys measurements, we found 30 of them could be moderately associated with genome-wide expression patterns, suggesting that gene regulation plays an important role in regulating neural responses to stimuli. Individual ephys features had distinct sets of genes associated with them; even for the same ephys feature, different interneuron subclasses had largely unique gene sets associated with that feature. We found that significant heritability of schizophrenia incidence is attributable to ephys loci, while we did not find significant heritability for any of the other psychiatric or neurological disorders tested. While our ephys response genes were surprisingly depleted of ion channels, we found an enrichment of adhesion genes in the ephys response gene set, consistent with adhesion molecules’ role in establishing neuron circuitry^50^ and distributing ion channels^51^. Finally, we observed that numerous adhesions genes exhibited functional switching between neuron subclasses; that is, they were mutually exclusively associated with distinct ephys functions in different neuron subclasses. Altogether, we propose that cell adhesion molecules play functionally distinct ephys roles in individual interneuron subclasses, and that ephys-associated genes play an important role in schizophrenia genetic risk.

## RESULTS

### RESULTS – Gene expression levels are predictive of diverse features of electrophysiological response to stimuli

We first set out to test the hypothesis that gene expression levels are predictive of ephys responses to stimuli. If true, this would suggest a significant role of transcriptional regulation in establishing a neuron’s response to stimulus. To test our hypothesis, we analyzed 3,120 paired measurements of gene expression levels and ephys responses of single interneurons drawn from four major GABAergic subclasses (Sst, Pvalb, Lamp5, Vip)^52^ (see Methods). Ephys responses for each neuron were summarized into a set of 52 ephys features (**Supplementary Fig. S1, Supplementary Table 1**), which we then predicted using a set of 12,986 gene expression levels measured in the same neurons. We focus on GABAergic neurons as they are one of the primary neuron types implicated in psychiatric disorders^53^.

We first trained statistical models to predict ephys response using all 3,120 interneurons across the four subclasses, and found that 30 of the 52 ephys features were well predicted (average Spearman rho > 0.5) (**Fig. 1a**), consistent with previous reports^52,54^. We hypothesized that the high predictive accuracy is driven by the fact that interneurons are globally distinct both transcriptionally and electrophysiologically^52^. Therefore, any gene that is differentially expressed between subclasses will also be predictive of ephys features that are different between subclasses, including subclass marker genes that are irrelevant to ephys response, for example. This is not desirable because genes such as markers are uninformative about ephys response biology.

**Figure 1:**
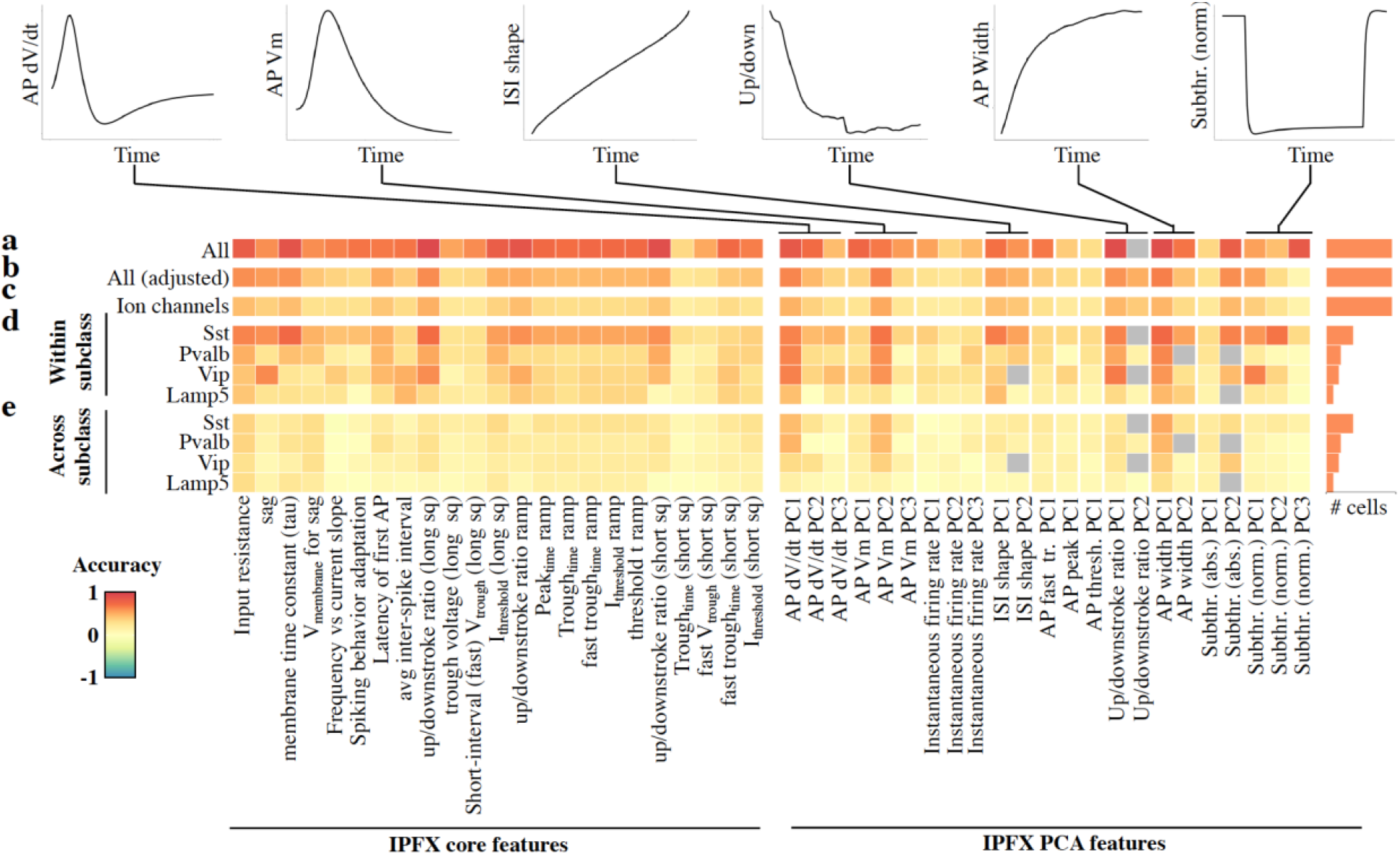
Transcriptional mechanisms of interneuron ephys response are subclass specific. (**a**) Average cross-validation accuracy of models predicting ephys response features using genome-wide gene expression levels, using data from all 3,412 interneurons across the four subclasses (Sst, Pvalb, Lamp5, Vip). Grey boxes indicate experiments on PCA-based features not executed because we computed fewer principal components. (**b**) Same as (a), but after removing average differences in gene expression and ephys features between subclasses before model training. (**c**) Same as (b), but models were only able to use the gene expression levels of 340 ion channels to predict ephys response features. (**d**) Same as (a), but now separate models are trained and validated for each of the four subclasses individually. (**e**) Average prediction accuracy of models trained on each subclass but used to predict ephys response of the other three subclasses.

We therefore removed average differences in gene expression levels and ephys features between subclasses before repeating model training on the same set of all 3,220 interneurons. By doing so, we force our predictive models to capture within-subclass covariation of gene expression and ephys response. The predictive performance of the models drop by an average of 32% per ephys feature (p < 3.61×10^−10^, Wilcox paired rank test) (**Fig. 1b**). These results confirm that the high predictive accuracy reported by others^52,54^ and in **Figure 1a** are driven by global differences in expression and ephys between interneuron subclasses, rather than true covariation of gene expression with ephys response. Still, we found 13 of 52 ephys features are well predicted (Spearman rho > 0.5) even after removing these global subclass differences in expression and ephys, suggesting that transcriptional regulation does play an important role in ephys response.

As a comparison group, we repeated our prediction experiments using a set of 340 annotated ion channels alone. Given the central role of ion channels in electrophysiology^55^, we reasoned our comparison would give quantitative insight into the extent to which variation in ephys response is explained by regulatory control of ion channels versus other mechanisms. On average, we found the ion channels-only model accuracy was 25.2% lower than the genome-wide model (p < 3.83 ×10^−10^, Wilcox paired rank test) (**Fig. 1c**), suggesting that there is significant unappreciated contribution of genes from outside of ion channel families towards ephys response in neurons that we will explore below.

### RESULTS – Electrophysiology response genes are subclass- and ephys feature-specific

Having established that many ephys features are well predicted by gene expression, we next wondered to what extent is the transcriptional ephys response subclass specific. It is already appreciated that different interneuron types have different ephys properties^52^ (**Supplementary Fig. S2**), but what has not been explored is whether the same genes underlie the ephys response of different interneuron classes. To test this hypothesis, we first constructed predictive ephys models separately for each of the four interneuron subclasses, and compared model prediction performance. When assessing the performance of each subclass-specific model on predicting ephys features for the same subclass on which they are trained, we observed wide variability in prediction performance for the same feature across subclasses (**Fig. 1d**). Among the four different subclasses, Sst ephys response features were the most predictive, with 17 of 52 features well predicted (Spearman rho > 0.5), while on average only four of 52 ephys features were well predicted for the other three subclasses. Surprisingly, for individual ephys features, there was large variation in prediction accuracy across the subclasses; only two ephys features (AP width PC1, AP dv/dt PC1) were common amongst the 10 best predicted ephys features per subclass, suggesting that the level of transcriptional regulation of ephys response differs substantially between subclasses. More specifically, Sst potentially has a clearer relationship between gene expression and ephys response compared to the other three subclasses. The ephys genes identified by the subclass-specific models also differed significantly from the all cells model – the average Jaccard index overlap between the subclass-specific models and the all cells model was only 2.9% (**Supplementary Fig. S3**), illustrating the importance of training subclass-specific models to identify ephys genes relevant to each subclass.

We further used each of our subclass-specific prediction models to predict the ephys responses of the other three interneuron classes (**Fig. 1e**). Every model performs significantly worse when predicting the ephys response of other subclasses compared to predictions of neurons within their own subclass (median decrease in accuracy of 52% across all 52 features and four subclasses, p < 1.55×10^−9^, Wilcox paired rank test). These results together suggest that even for individual features of neuron ephys response, interneuron classes exhibit significantly different transcriptional ephys responses.

While our results above demonstrate high specificity of each subclass’s transcriptional response to ephys response, they do not directly measure the extent of overlap of the genes underlying these transcriptional responses. We therefore next tested two more refined hypotheses. First, we hypothesized that within the same interneuron subclass, different genes contribute to different ephys response features. Second, we hypothesized that even for the same ephys feature, different genes contribute to the response for different interneuron subclasses.

To test these two hypotheses while avoiding potential feature selection bias in our predictive models (see **Methods**), we identified the top 100 genes most correlated with each of the 42 common ephys features measured in all four interneuron subclasses, termed the correlation-ephys genes. First, for each subclass individually, we calculated the number of times a given gene was identified as a correlation-ephys gene for different ephys features. For each subclass then, each ephys feature was assigned an ephys feature specificity score, defined as the percentage of correlation-ephys genes shared with fewer than five ephys features in the entire subclass. On average across the 42 ephys features in Sst neurons for example, 25% of correlation-ephys genes were only correlated to less than five ephys features, indicating a high degree of specificity of genes to specific ephys features (**Fig. 2a**); other subclasses showed similar levels of specificity (**Supplementary Fig. S4**). These results indicate different genes contribute to different ephys response features in the same subclass.

**Figure 2:**
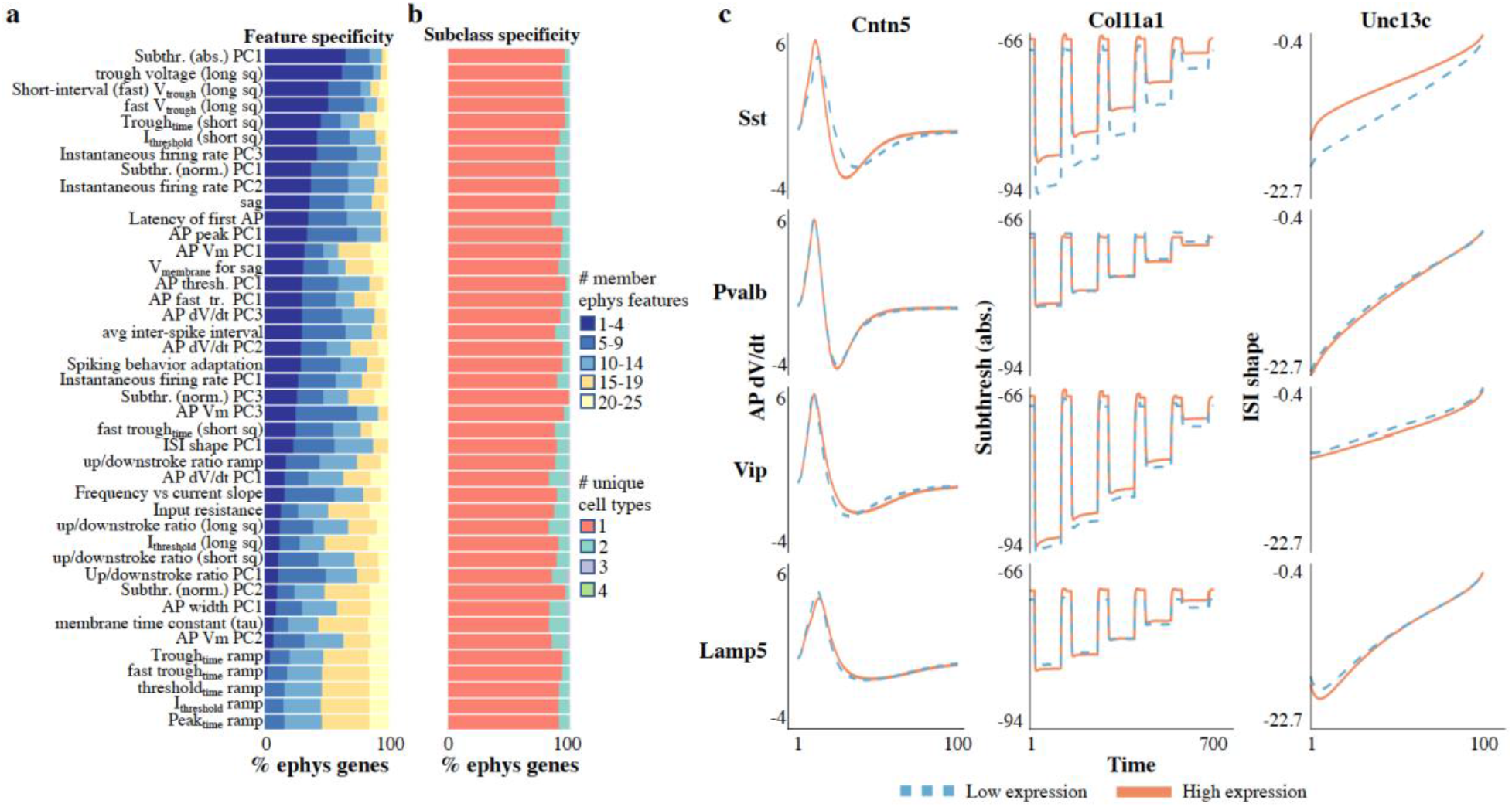
Ephys response genes are subclass- and ephys feature-specific. (**a**) Stacked bar chart indicating, for each Sst ephys feature, the fraction of the corresponding correlation-ephys genes that are also correlation-ephys genes for other ephys features in Sst. Correlation-ephys genes are grouped based on how many other ephys features they are correlated with. Features are sorted based on average number of other ephys features that each ephys gene is correlated with. (**b**) Stacked bar chart indicating, for each ephys feature, the percentage of all genes correlated with that feature (taken over all four subclasses) that are shared by either one (unique), two or three subclasses. No ephys gene was correlated with the same ephys feature across all four subclasses. (**c**) Plot of average AP dV/dt, Subthresh. (abs.), and ISI shape as a function of time steps for the genes Cntn5, Col11a1 and Unc13c, respectively. For AP dV/dt, time point 1 references the initiation of the first action potential. For Subthresh. (abs.), values represent the concatenated responses to hyperpolarizing current steps (from -10 pA to -90 pA, in steps of 20pA). For ISI shape, the average of ISI voltage trajectories are visualized, aligned to the threshold of the initial AP and normalized in duration. Each of the three genes are ephys genes specific to Sst, and are not ephys genes of the other three subclasses. Lines indicate the average ephys behavior of neurons at the top (orange) and bottom (blue) 10^th^ percentile in expression of the three genes for each of the four subclasses.

Next, to establish the extent of gene overlap of ephys genes for the same ephys feature across subclasses, we calculated a subclass specificity score measured as the average percentage of correlation-ephys genes specific to each subclass, when considering the same ephys feature across subclasses. We found on average subclass specificity of 90.6% across the 42 ephys features (**Fig. 2b**), indicating that even for the same ephys feature, different genes establish a response to ephys stimulus. Our results were consistent when we performed this analysis on the genes identified by our predictive models: we found minimal overlap in genes selected by models trained on the same interneuron class but across different ephys features (**Supplementary Fig. S5**), as well as minimal overlap in genes selected by models trained on the same ephys feature but across interneuron classes (**Supplementary Fig. S6**) (median Jaccard index < 0.92%). In total, these results suggest that each ephys feature is associated with a distinct set of genes within a given subclass, and even for the same ephys feature, the sets of genes associated with that feature are distinct for each interneuron type.

To visualize the subclass-specific effect of ephys genes, **Figure 2c** illustrates the first action potential recorded for neurons from each subclass, when averaged for neurons exhibiting either low or high levels of expression of the three ephys genes Cntn5, Cdh7 and Reln predicted to specific to Sst. For each of the three ephys genes, a significantly larger difference in action potential behavior is observed for Sst interneurons. Reln, one of the most positively correlated genes associated with action potential’s upstroke downstroke ratio in Sst (Pearson rho = 0.34, rank 1), showed a much higher upstroke downstroke ratio in Sst cells when strongly expressed under long square current stimulation, suggesting a sharper spiking pattern. Importantly, Reln is negatively correlated (Pearson rho = -0.21, rank 55) in Lamp5, and is not an ephys gene for Pvalb or Vip (**Supplementary Fig. S7**), suggesting Reln is functional in only Sst and Lamp5 cells. Reln, an extracellular matrix protein with a primary role in synaptic plasticity, is particularly of interest due to its implication in psychiatric disease risk with evidence from animal models, GWAS, epigenomics, expression analysis spanning across multiple ethnicities^56–59^. Our finding strongly suggests that Reln could be influencing single cell neuronal action potential in a subclass specific manner, complementing previous work showing deletion of the human homolog RELN in iPSC-derived neurons leads to changes in the dendritic morphology and synaptic components^59^ of neurons. Our results therefore suggest that the genes necessary to explain ephys variation are different between subclasses as well as ephys features, revealing a distinct gene ephys response program between interneurons and ephys features.

### RESULTS - Significant Schizophrenia heritability in Sst action potential voltage change genes

Having identified genes associated with multiple features of ephys responses and interneuron subclasses, we next tested our hypothesis that significant psychiatric disorder heritability could be attributed to one or more ephys response features. Previous partitioned heritability analyses of psychiatric disorders focused on identifying a general role for neurons and interneurons^15,53^ as a class of target cells of psychiatric genetic variants, but thus far have not explored the potential role of subclass-specific genes associated with single neuron ephys response.

We first performed partitioned heritability analysis to confirm that interneurons are a strong candidate target subclass for psychiatric variants (**Fig. 3a**). We found that Sst and Pvalb interneurons are implicated specifically in Schizophrenia and bipolar disorder (**Fig. 3a**), consistent with previous work^60^. Given that we observed Sst had the strongest association between gene expression and ephys responses (**Fig. 1d**), we focused on Sst cells for the remainder of this section.

**Figure 3:**
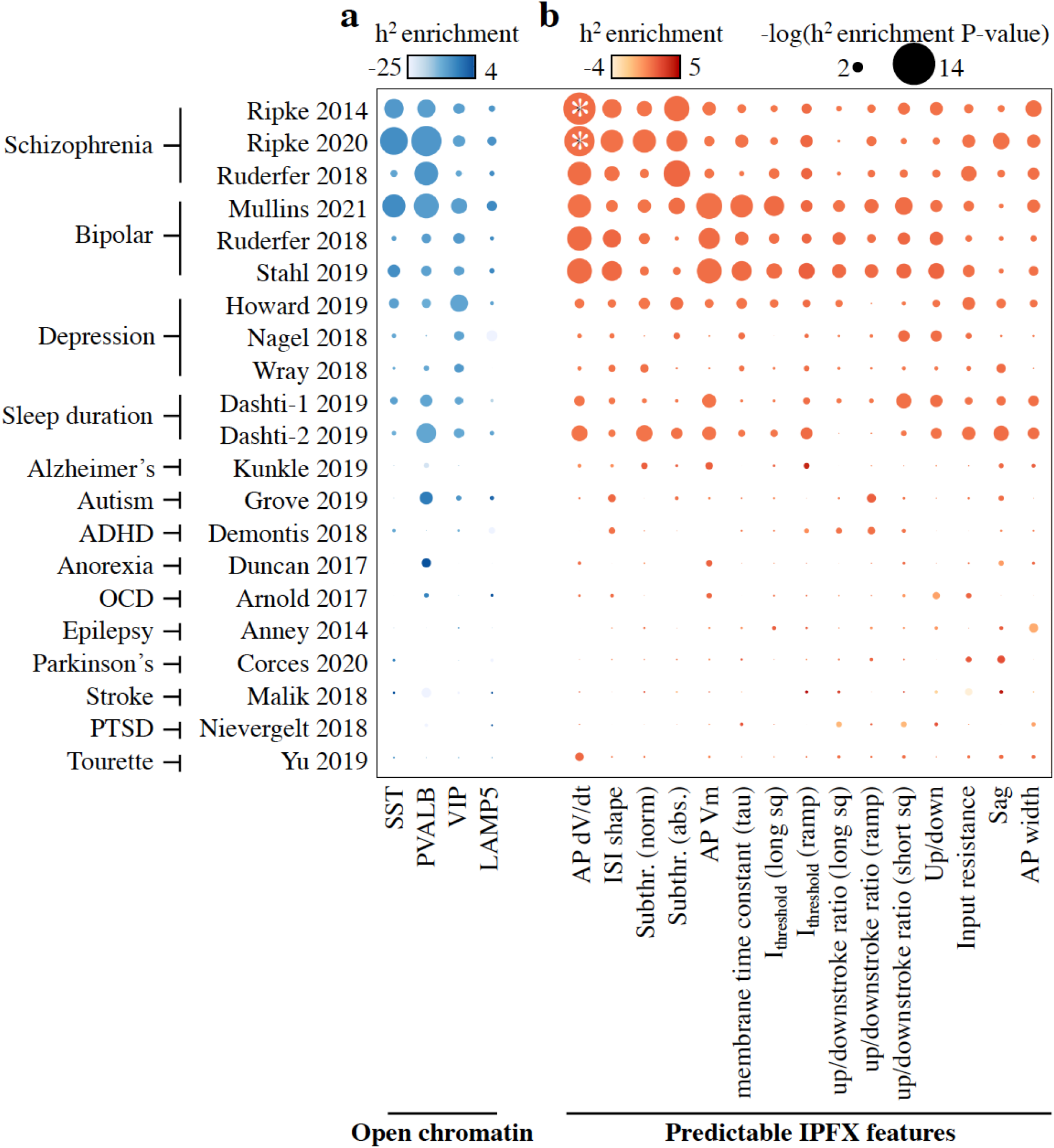
Significant Schizophrenia heritability is attributable to first action potential voltage change genes in Sst neurons. (**a**) Plot of heritability analysis of GWAS traits (rows) with respect to subclass-specific open chromatin regions in human. Circle size represents the magnitude of h^2^ p-values, while circle color intensity represents the h^2^ enrichment from LD score regression. GWAS datasets are named with the first author and year of publication, organized by trait type. (**b**) Same as (a), except heritability analysis is with respect to different ephys response gene sets inferred for Sst neurons. White asterisks indicate significant p-value (p < 0.05) after Bonferroni correction of permutation-based p-values.

For each of the 15 IPFX core and PCA ephys features well predicted in Sst cells (Spearman rho > 0.5 and 0.4 respectively; **Fig. 1d**), we extracted the genes associated with that feature from our predictive models. We then mapped them to their human ortholog, and used LD score regression^61^ to identify significant heritability of psychiatric disorders with respect to the ephys response gene set. We found strong Schizophrenia heritability with respect to the first action potential voltage change (AP dV/dt) ephys genes (nominal p = 1.14 × 10^−10^, permuted p < 0.001) (**Fig. 3b**), which replicated across multiple SZ GWAS,and passed Bonferroni correction of both the nominal and permutation p-value. The heritability signal of the AP dV/dt feature was specific to Schizophrenia; no other psychiatric disorders or neural disorders showed significant heritability, though Bipolar disorder shows more nominal signal than other traits (**Fig. 3b**). The specificity of the heritability signal to Schizophrenia does not appear to be driven by statistical power of the underlying GWAS; there were other tested disorders that had substantially larger cohort sizes and number of genome-wide significant loci in their GWAS (**Supplementary Table 2**). These results suggest that action potential voltage changes in Sst ephys responses may play a role in mediating Schizophrenia genetic risk.

To shed insight into the molecular mechanisms by which the Sst AP dV/dt ephys genes influence action potential voltage changes, we first reasoned that the Sst AP dV/dt ephys gene set likely contained genes not causally responsible for ephys response, but were only indirectly correlated. Given that only 2% of the Sst AP dV/dt ephys genes were annotated ion channels (median percentage of Sst ephys gene sets that were ion channels is 4.61%; **Supplementary Table 3**), we reasoned that causal ephys genes would be more likely to be also associated with ion channels. We therefore inferred a gene co-expression network of the Sst AP dV/dt genes with ion channels, and visualized their interactions (**Fig. 4a**). Surprisingly, of the 676 AP dV/dt genes, 93 of them form a single, connected module of strongly co-expressed genes (**Fig. 4a**), while most remaining genes were isolated nodes in the network (**Supplementary Fig S8**). Compared to the other connected gene modules from the 15 well predicted ephys features from **Figure 3b**, this 93-gene AP dV/dt gene module is enriched in cell cell adhesion, synapse signalling, neuronal morphology, membrane components, and nervous system development pathways (**Fig. 4b**), with cell adhesion showing the strongest enrichment (p < 8.32 × 10^−3^, FDR corrected GSEA test). Within the module includes Rgs6, a genome-wide significant SZ locus who directly regulates (GIRK)-potassium channel activation via the inhibition of GABAb receptors^62^, in addition to covarying strongly in expression with ion channels Grin2a, Grin3a and Cacna2d3. Interestingly, Rgs6 also co-varies with the gene Reln (**Fig. 4a**), a known SZ associated gene^56^ that promotes proper neuronal positioning via adhesion as well as dendritic development^63,64^. These results suggest the AP dV/dt gene module is likely to play a functional role in ephys response, possibly through cell adhesion.

**Figure 4:**
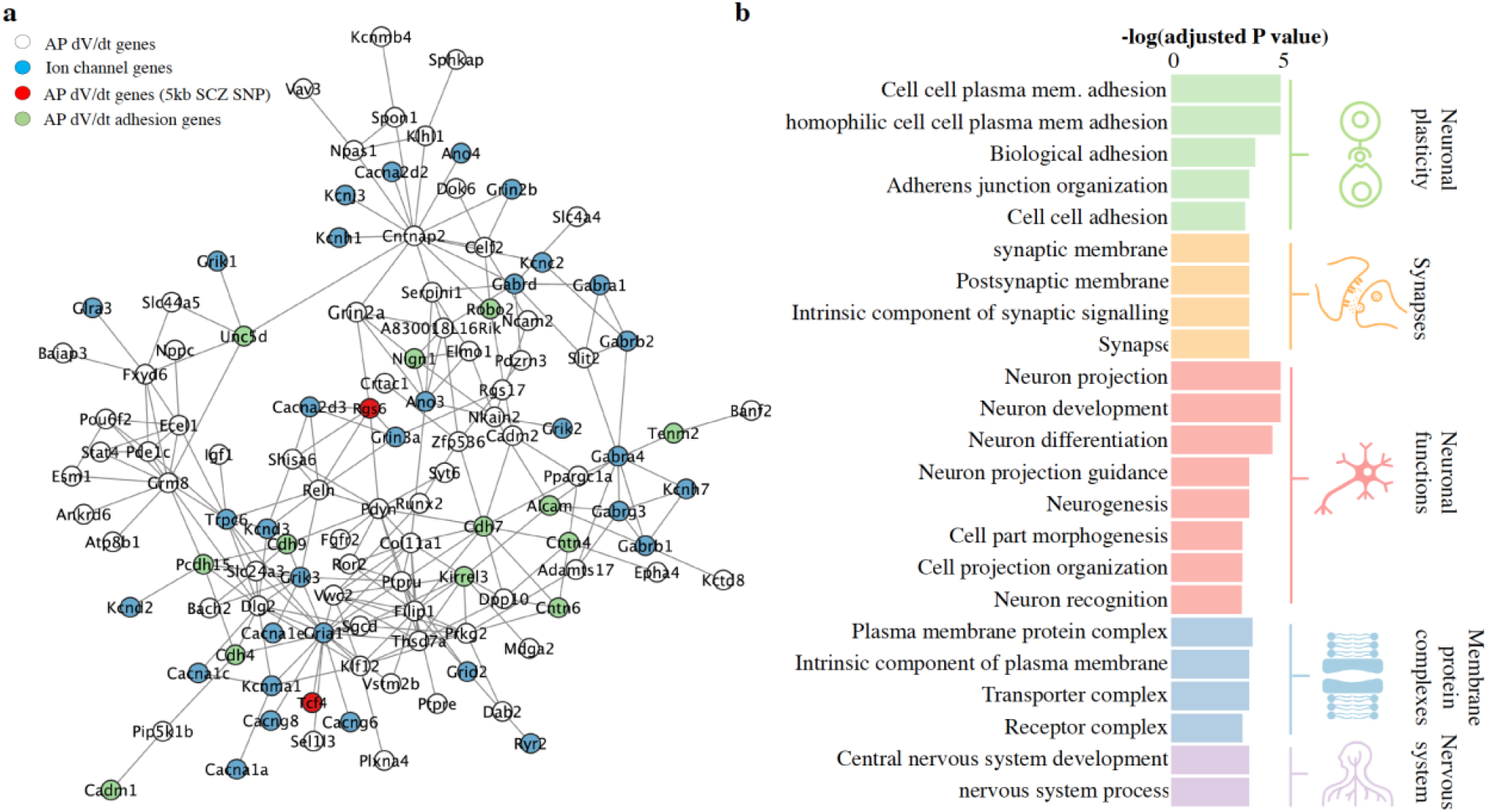
First action potential voltage change genes form a co-expression module enriched in cell adhesion genes. (**a**) Visualization of the Sst first AP dV/dt ephys response gene co-expression network, including correlated ion channels. Isolated first AP dV/dt genes not connected to any other genes were dropped; the full network figure is available in **Supplementary Fig. S8**. White nodes represent AP dV/dt genes, green nodes represent the subset of AP dV/dt genes also annotated as adhesion genes (GO:0098742), red nodes indicate AP dV/dt genes also within 5kb of a SCZ GWAS SNP, and blue nodes represent ion channels. (**b**) Gene set enrichment analysis of first AP dV/dt gene module from (a), with respect to all AP dV/dt genes including the isolated nodes. All enrichments shown were significant (q < 0.05) after FDR correction.

### RESULTS – Cell adhesion ephys genes are versatile and can switch roles between subclasses

Given the strongest enrichment for Sst first AP dV/dt genes was cell adhesion as well as the surprisingly depleted representation of ion channels, we wondered whether cell adhesion genes played a more general role in other ephys features as well. Across the other 15 well predicted ephys features in Sst, we found cell adhesion genes were enriched in four ephys feature sets, composed of AP dV/dt, AP width, and up/downstroke ratios (**Supplementary Fig. S9**). The enrichment of cell adhesion genes across multiple Sst ephys features was surprising, given we saw the poor overlap in associated genes between different ephys features of Sst (**Fig. 2a**), and poor overlap of ephys features between cell types (**Fig. 2b**).

To obtain a global view of the roles of cell adhesion in ephys, for each cell adhesion gene, we identified the subset of ephys features in each interneuron subclass for which the cell adhesion gene was also a correlation-ephys gene as defined above. We also grouped the 42 ephys features into six distinct ephys function categories: AP peaking features, AP threshold features, hyperpolarization features, AP train features, AP trough features and electrical capacity features (**Supplementary Table 1**). We observed that 237 cell adhesion genes were also correlation-ephys genes across all subclasses. Surprisingly, of these 237 cell adhesion ephys genes, all of them were correlated with different ephys features in different subclasses.

Kirrel3 in particular stood out because it is correlated with 19 distinct ephys features across all subclasses, was correlated with different ephys features in different subclasses (**Supplementary Fig. S10**), and maintained high and consistent expression across all subclasses (**Supplementary Fig. S11**). **Figure 5a-c** illustrates how Kirrel3 is highly correlated with the frequency vs current slope ephys feature in Lamp5 but not Pvalb. In contrast, **Figure 5d-f** shows that Kirrel3 is the highly correlated with the AP dV/dt PC1 ephys feature in Pvalb, but not Lamp5. These differences in ephys feature correlations across subclasses are not explained by changes in mean expression level; Kirrel3 is not differentially expressed in any subclass. To determine how Kirrel3 may be changing its ephys correlation patterns in different subclasses, we visualized the local gene co-expression network of Kirrel3 in both Pvalb and Lamp5 (**Fig. 5g,h**); a global co-expression network including Kirrel3 is visualized in **Supplementary Fig. S12**. In Pvalb neurons, Kirrel3 co-expresses with mostly AP dV/dt ephys genes, whereas in Lamp5 neurons, Kirrel3 co-expresses with mostly firing frequency vs current slope ephys genes. **Figure 5i** summarizes the function categories of features that Kirrel3 is most correlated with, and illustrates how Kirrel3 is correlated with a significant number of ephys features in several function categories, but the specific categories differ across the subclasses. These results in total suggest that Kirrel3 may be broadly orchestrating “switching” of neuronal ephys functions in distinct subclasses by co-expressing and interacting with distinct sets of gene modules.

**Figure 5:**
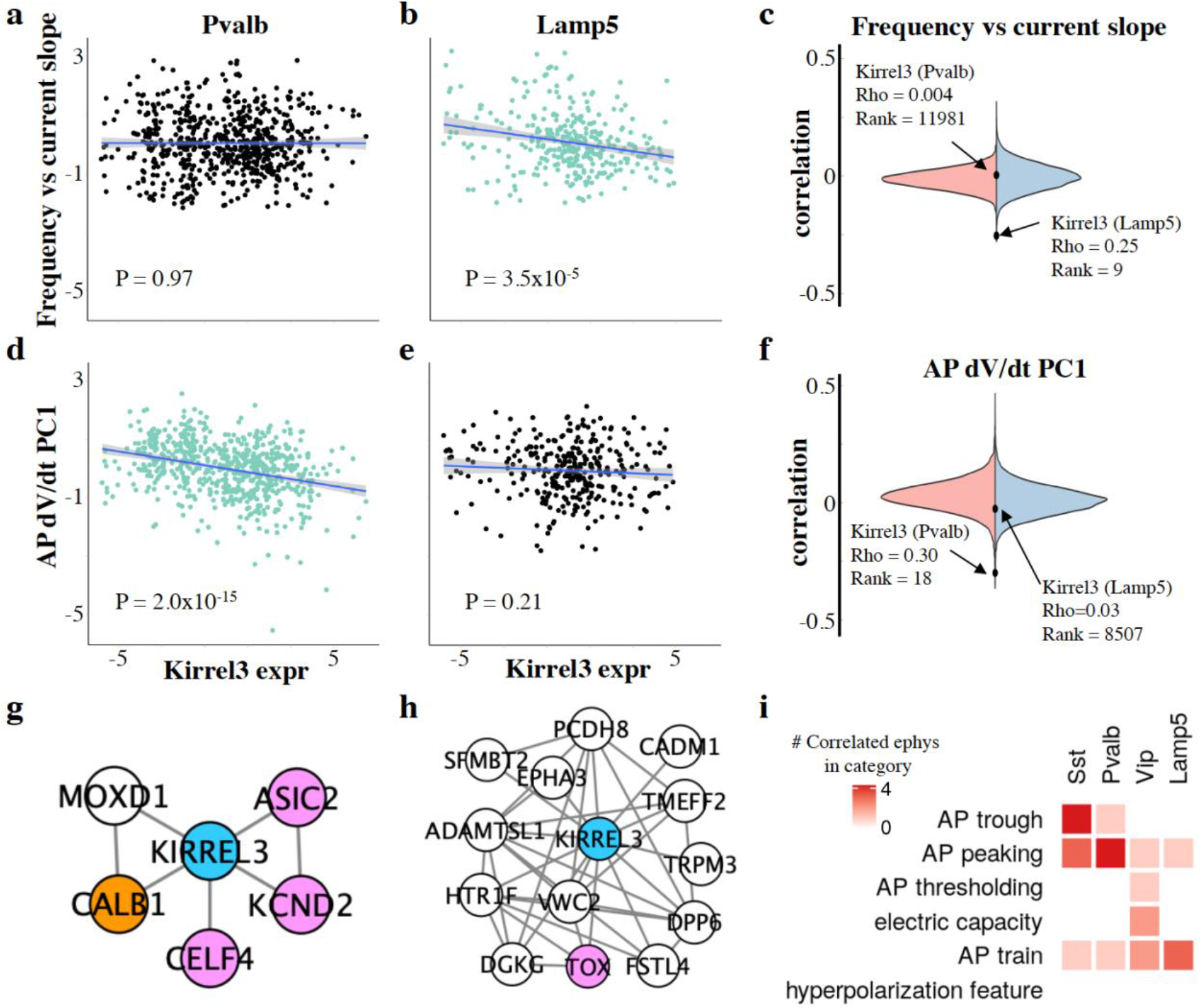
Cell adhesion genes exhibit function switching across interneuron subclasses. (**a**) The estimated frequency vs current slope ephys feature as a function of Kirrel3 expression for Pvalb neurons. (**b**) Same as (a), but for Lamp5 neurons. (**c**) Distribution of correlations of all **12986** genes with the Frequency vs current slope ephys feature for both Pvalb (left) and Lamp5 (right). Shown as closed circles are the correlations of Kirrel3 with frequency vs current slope in both cell types, as well as the rank of Kirrel3 when genes are sorted by decreasing magnitude of correlation. (**d**) The AP dV/dt PC1 ephys feature as a function of Kirrel3 expression for Pvalb neurons. (**e**) Same as (d), but for Lamp5 neurons. (**f**) Distribution of correlations of all **12986** genes with the AP dV/dt ephys feature for both Pvalb (left) and Lamp5 (right). Shown as closed circles are the correlations of Kirrel3 with frequency vs current slope in both cell types, as well as the rank of Kirrel3 when genes are sorted by decreasing magnitude of correlation. (**g**) Local co-expression network of Kirrel3 in Pvalb neurons. Pink nodes represent correlation-ephys genes associated with AP dV/dt PC1 in Pvalb, white nodes represent correlation-ephys genes associated with Frequency vs current slope in Pvalb, and orange nodes are correlation-ephys genes associated with both ephys features. (**h**) Local co-expression network of Kirrel3 in Lamp5 neurons. Node coloring is the same as in (g). (**i**) Heatmap indicating the relative number of ephys features in each category and subclass that Kirrel3 is correlated with.

Finally, we next determined the extent to which other cell adhesion ephys genes exhibit ephys function switching. We first identified a set of 83 high confidence cell adhesion ephys genes as those cell adhesion genes that are correlated with more than 6 ephys features in total. Of the 83 high confidence cell adhesion ephys genes, 34 of them were identified as exhibiting switching behavior between at least two subclasses, including Kirrel3 (see **Methods**). Furthermore, of these 34 genes, 19 of them show clear one to one binary switching patterns in which a gene is exclusively associated with one ephys function category in one subclass, and exclusively associated with another ephys function category in another subclass (**see Methods**). This finding strongly suggest that adhesion molecules more generally could be orchestrating different set of ephys functions between different subclasses, leading to diversity of neuronal electrical properties.

## DISCUSSION

One of our surprising observations was the very low overlap between the ephys response genes and known ion channels, given the central role of ion channels in electrophysiology^65^. Still, our ephys response genes were systematically better predictors of ephys response than ion channels (**Fig. 1**). There are several possible explanations. First, some ephys response gene sets are enriched in functions such as cell adhesion that have established roles in regulating surface expression and activity of ion channels, although they do not directly transport ions themselves. Several genes, including some cell adhesion genes, are also correlated with ion channel genes, suggesting some shared functions. Perhaps ion channels are critical for establishing baseline ephys response, but other genes may be responsible for modulating the behavior of those responses. Second, identification of ephys response genes involved first using the IPFX software package to extract quantitative features from patch clamp experiments; it is possible that ion channels may be more responsible for variation in ephys reponse for features that have not yet been extracted.

Our work identified a significant role of cell adhesion genes in the AP dV/dt ephys response feature, as well as correlation of their expression with ion channels. The result is consistent with the established role of adhesion molecules in directly and mechanistically determining neuronal excitability and the firing event in neurons, through direct interaction with ion channels to regulate their properties, surface expression and patterning. Enrichment of cell adhesion genes is specific to a subset of ephys features including AP dV/dt, Up/downstroke ratio, AP width, sag, and I_threshold_: features such as interspike intervals shows no such significant enrichment in adhesion pathways but is instead more enriched in synaptic components. Importantly, we observed the same enrichments in adhesion molecules in the correlation-ephys genes identified independently of our GLM models.

Another interesting observation we made was the extensive ephys function switching across subclasses by nearly half of all high confidence, cell adhesion ephys genes (30/61). As far as we are aware, this function switching has not previously been reported. Our co-expression network analysis supports the idea that the same cell adhesion gene can interact with systematically different sets of genes (including other cell adhesion genes) in different subclasses even without changing significantly in gene expression levels, suggesting they are being activated as parat of different pathways in different subclasses. Further investigation knocking down these switching adhesion molecules in different subclasses to probe the resulting downstream change in ephys patterns is warranted.

Our work identified significant heritability of AP dV/dt in SCZ, the first reported as far as we know. Furthermore, the genes responsible for the heritability signal show significant enrichment in adhesion molecules. There are some caveats with our heritability analysis. Our partitioned heritability analysis of Schizophrenia depended on identification of ephys response genes via statistical analysis of paired gene expression-ephys neuron measurements. As such, it is possible that some of the ephys response genes we identified are false-positive ephys response genes, in that they may not be directly involved in ephys response but are correlated with genes that are directly involved. To prevent our partitioned heritability analysis from over-estimating contributions of ephys response genes to our heritability estimates, we based our conclusions on a stringent permutation test which explicitly controlled for the number of ephys response genes identified, as well as the gene-gene correlation structure in each interneuron class. Our criteria is likely overconservative, and underestimates the contribution of single neuron ephys in psychiatric genetics. Note that the gold standard approach to ephys gene identification would in principle involve ephys characterization before and after gene perturbations of individual interneurons. However, given the prohibitive labor and cost of ephys phenotyping for such large screens, association analyses such as ours are useful for prioritizing individual candidate ephys response genes.

In our study, we identified ephys response genes in mouse before mapping them to their human orthologs in order to perform heritability analysis on mental disorders. Our analysis in mouse was motivated by the fact that the largest publicly available dataset in humans was an order of magnitude smaller^66^, it only profiled excitatory neurons, and we observed poor overall association between gene expression levels and ephys response (**Supplementary Fig. 13-14**). While human and mouse orthologs do not necessarily share similar function, we restricted ourselves to only considering genes with 1-1 orthologs in human. Such unambiguous orthologs are more likely to maintain similar gene function compared to other genes^67^.

## ACKNOWLEDGEMENTS

GQ was supported by NSF CAREER award 1846559. This work was funded in part by grants from the National Institutes of Health (NIH), including the Office of the Director/National Institute of Mental Health (DP2 MH129987, GQ), and the National Institute of Child Health and Human Development (P50 HD103526).

## DATA AVAILABILITY

This is a computational study and no new data were generated.

## PREVIOUSLY PUBLISHED DATASETS

Integrated Morphoelectric and Transcriptomic Classification of Cortical GABAergic Cells: Nathan W et al, 2020, https://portal.brain-map.org/explore/classes/multimodal-characterization, Author’s website; Human neocortical expansion involves glutamatergic neuron diversification: Jim Berg et al, 2021, https://portal.brain-map.org/explore/classes/multimodal-characterization, Author’s website; Single-cell epigenomic analyses implicate candidate causal variants at inherited risk loci for Alzheimer’s and Parkinson’s diseases: M. Ryan Corces et al, 2020, https://www.nature.com/articles/s41588-020-00721-x, Supplementary Table 7

## METHODS

### GWAS summary statistics acquisition

We obtained GWAS summary statistics for the following phenotypes: Tourette Syndrome^14^, obsessive compulsive disorder^12^, schizophrenia^2,3,68^, bipolar disease^3–5^, autism spectrum disorder^9^, attention-deficit/hyperactivity disorder^10^, post traumatic stress disorder^13^, major depression disorder^6^, anorexia, autism, epilepsy, multiple sclerosis, parkinson disease, sleep duration, stroke and alzheimer’s disease. The full GWAS manifest that documents the date of the download, URL link to the download locations as well as the file names of the GWAS are documented in the **Supplementary Table 2**.

### GWAS summary statistics preprocessing

All GWAS summary statistics were processed using an LDscore^69^ GWAS preprocessing pipeline (https://github.com/bulik/ldsc/wiki/Heritability-and-Genetic-Correlation), with the exception of the summary statistics for major depression (PGC_UKB_depression_genome-wide.txt) for which we first exponentiated the log odd ratio. Briefly, the pipeline involved filtering the SNPs to the HapMap3 SNPs (downloaded from https://alkesgroup.broadinstitute.org/LDSCORE/, under the file name w_hm3.snplist on August 2020), followed by the removal of strand ambiguous SNPs that yield errors in the downstream partitioned heritability analysis.

### Human-mouse ortholog mapping and ion channel annotations

Ortholog mappings between the human and mouse genome was downloaded from Biomart on February 9, 2021, using the GRCh38 build (Ensembl 102). Human-mouse gene pairs were filtered to those labeled as ‘high confidence’ orthologs by Ensembl. We then only retained 1-1 human-mouse pairs. Gene coordinates in the GRCh37 build were obtained using the biomaRt^70^ R package on July 9, 2021. In addition to the human mouse ortholog mapping list, we also downloaded a curated list of ion channel genes from https://www.genenames.org/data/genegroup/#!/group/177.

### Human and mouse Patch-seq data acquisition

Mouse Patch-seq^52^ transcriptomics data and accompanying meta data for the L2/3 visual cortex GABAergic neurons were obtained on April 2021 from https://portal.brain-map.org/explore/classes/multimodal-characterization/multimodal-characterization-mouse-visual-cortex, under the file name 20200513_Mouse_PatchSeq_Release_count.v2.csv.tar and 20200711_patchseq_metadata_mouse.csv. Accompanying electrophysiology feature measurements stored in NWB files generated by Allen Institute for Brain Science (https://portal.brain-map.org/explore/classes/multimodal-characterization) were acquired via the Distributed Archives for Neurophysiology Data Integration (DANDI, https://gui.dandiarchive.org/#/dandiset, ID:000021, download date: August 2021).

Human Patch-seq^66^ transcriptomics and metadata for excitatory neurons were downloaded from BRAIN/NeMo using Aspera on Dec 2020, while the corresponding electrophysiological measurements stored in NWB files were acquired via the Distributed Archives for Neurophysiology Data Integration (DANDI, https://gui.dandiarchive.org/#/dandiset, ID:000023, download date: September 2020).

### Within subclass and across subclass Patch-seq RNA preprocessing

**M**ouse Patch-seq RNA data was first subsetted to the Sst, Pvalb, Sncg, Vip, and Lamp5 cells to keep the most represented subclasses. We further removed cells with the aspiny dendrite type as well as the Sst Chodl sub-subclass due to their substantial difference in transcriptomics profiles from the remaining Sst cells. We clustered the cells using their RNA measurements and removed clusters with a heterogeneous mixture of subclasses (**Supplementary Fig. S15**, cluster 4 and 7). We further filtered out cells with low count and low gene number (< 200,000 counts and < 4,000 genes expressed), followed by removal of genes expressed in fewer than 30 cells, yielding the mouse **Patch-seq QCed count matrix**. We then log normalized the mouse Patch-seq QCed count matrix with a scaling factor of 10^6^, then subsetted to high confidence human-mouse 1-to-1 ortholog genes. We used a linear model to regress out age, biological sex, major subclass, transcriptomics batch, hemisphere and structure effects. For transcriptome batches and structure groups that were small (< 15 cells), we grouped all small batches together (within each annotation separately) before batch correction. Both the “within subclass RNA processing” and “across subclass RNA processing” procedures are exactly as described above.

With respect to the human data, human Patch-seq RNA data was first filtered for missing information on structure and medical conditions. From the remaining cells, we then filtered for cells with extremely high count number and genes expressed (> 2 million counts, 20, 000 genes expressed) as well as cells with low gene expression (< 5000 genes expressed). We then removed the genes expressed in less than 3 cells, yielding what we refer to as the human Patch-seq QCed count matrix. We then log normalized the human Patch-seq QCed count matrix with a scaling factor of 10^6^, then subsetted to high confidence human-mouse 1-to-1 ortholog genes. We used a linear model to regress out age, biological sex, major subclass, medical conditions, hemisphere and structure effects. Due to the small sample size in all subclasses except FREM3 cells (average < 34 cells per subclass other than FREM3), we only conducted the within subclass analysis using FREM3 cells.

### All cells Patch-seq RNA preprocessing

The All cells Patch-seq RNA processing procedure is the same as the within subclass Patch-seq RNA preprocessing procedure, with the exception of the batch correction step where we do not regress out the major subclass, transcriptomics batch and structure due to their partial correlation with subclass label. The “All (adjusted) Patch-seq RNA processing” procedure is the same as the All subclass preprocessing procedure, with the exception of also correcting for the subclass label in addition to the other covariates (age, biological sex and hemisphere).

### Electrophysiological raw feature extraction

We used the Allen SDK package (v0.16.3, https://github.com/AllenInstitute/AllenSDK) to import the electrophysiological data stored in the NWB files. We used the IPFX package (v1.0.4, https://ipfx.readthedocs.io) to extract two different sets of electrophysiological features, which we have termed the “IPFX core features” and “IPFX feature vectors”. The sweeps tagged with failure messages were removed by the built-in quality control (QC) *drop_failed_sweeps* step and 47 IPFX core features were extracted using the *extract_data_set_features* function. We extracted the IPFX feature vectors using the *run_feature_vector_extraction* function on the same set of cells that remained after subclass-specific Patch-seq RNA preprocessing, which produced 11 “intermediate” feature vectors. These 11 intermediate feature vectors include single-neuronal responses to hyper-polarizing current steps, action potential waveform, inter-spike-interval shape, instantaneous firing rate across long steps, together with upstroke/downstroke ratio, peak, fast trough, threshold, and width at half-height across long steps^71^ (including action potential waveform, subthreshold response waveform among others). The 11 intermediate feature vectors capture the responses of each single cell to three stimulus types: short-square (3ms), long-square (1s), and ramp current pulses stimuli, which has been described in previous publications of multimodal characterization of individual neurons using Patch-seq^52,66,71^. We excluded the ramp-specific segments of the “first_ap_v” and “first_ap_dv” feature vectors as was done previously^71^ due to a number of neurons not spiking under ramp stimulation. Since most cells were only stimulated under the +0pA, +40pA and +80pA rheobase sweep, we further retained only the segments corresponding to these sweeps for downstream analysis for “inst_freq”, “spiking_fast_trough_v”, “spiking_peak_v”, “spiking_threshold_v”, “spiking_width”, and “spiking_upstroke_downstroke_ratio” feature vectors. **Supplementary Table 1** lists all IPFX features along with a short description of each, and also indicates which intermediate feature vectors were reduced.

### Within subclass IPFX core feature preprocessing

From the raw IPFX core features extracted above, we subsetted to the common set of cells between the raw IPFX core features and the mouse Patch-seq QCed count matrix. We then removed IPFX core features with more than 750 missing values, as well as cells with more than ten missing IPFX core features. For each of the subclasses individually, we deemed the top and bottom 1% of values for each IPFX core feature as outlier values and set them to NA. We then computed z-scores for each feature, and corrected for age, biological sex, major subclass, transcriptomics batch, hemisphere and structure using a linear model followed by a second round of outlier removal, again defined by the top and bottom 1% of values for each IPFX core feature. Finally, we z-score each of the ephys features within each of the major subclasses.

### Within subclass IPFX PCA feature vector preprocessing

From the 11 intermediate feature vectors extracted above, we subsetted the raw feature data to the same set of cells as the mouse Patch-seq QCed count matrix. We then z-scored each of the 11 intermediate feature vectors, and for each intermediate feature vector, we corrected for age, biological sex, major subclass, transcriptomics batch, hemisphere and structure using a linear model. For each subclass separately, we then z-scored the residuals and extracted the fewest number of PCs that explained at least 90% variance for each of the 11 intermediate feature vectors. These PCs then formed the 11 IPFX PCA feature vectors.

### All cells IPFX core and PCA feature preprocessing

The All cells IPFX core feature preprocessing is the same as the within subclass IPFX core feature preprocessing, with the exception that during batch correction, we only correct for age, biological sex and hemisphere. To perform All cells IPFX PCA feature vector preprocessing, from the IPFX intermediate features, we subsetted the raw feature data to the common set of cells in both the raw IPFX core features and the mouse Patch-seq QCed count matrix. We then z-scored each of the 11 IPFX intermediate feature vectors separately and corrected for age, biological sex and hemisphere. We further scaled the residuals and computed PCs using all the remaining cells from all four subclasses.

For the “All (adjusted) cells IPFX core and PCA feature preprocessing”, we used the same procedure as the All cells preprocessing procedure with the exception of also correcting for the subclass label along with the rest of the covariates (age, biological sex and hemisphere).

### Across subclass IPFX core and PCA feature vector preprocessing

Across subclass IPFX core feature preprocessing is the same as the Within subclass IPFX core feature preprocessing. Across subclass IPFX PCA feature preprocessing is the same as the All cells IPFX PCA feature preprocessing, with the exception of the calculation of PCA. In our across subclass model evaluation (**Fig. 1e**), we are using models trained using a single subclass to predict ephys response in the other three subclasses. Therefore, during model training, we compute PCs for each subclass separately. During model evaluation, the IPFX intermediate feature vectors of the test subclasses are projected onto the PCs of the subclass that the model is trained on.

### Prediction of electrophysiological features from gene expression patterns (all cells, within subclass, ion channel only)

We performed three sets of ephys predictions based on gene expression patterns: all cells, within subclass and across subclass. For the all cells and within subclass prediction experiments, we followed the same general prediction and evaluation strategy, but only switched the input dataset. We employed 3-fold nested cross validation, repeated for five different seeds and averaged to estimate prediction accuracies. For each seed and fold of the outer cross validation, the training data was used to train 10 elastic net-regularized GLMs with alphas of zero to one with 0.1 increments implemented using cv.glm function from the glmnet package^72^ in R. From these 10 GLMs, we identified the GLM with highest cross validation accuracy on the training data, and used it to predict the test fold’s electrophysiology given the expression data. Test accuracy was measured as Spearman correlation between the predicted electrophysiology and the measured electrophysiology of the test fold, then averaged across all three test folds for all five seeds to get an average Spearman rho per ephys feature. For the all cells experiments, the input data to this evaluation procedure was the set of all mouse interneurons across the four subclasses (Sst, Pvalb, Vip, Lamp5), not corrected for average subclass differences in expression or ephys features (All cells Patch-seq RNA preprocessing section). For the within subclass experiments, this evaluation procedure was applied to each of the four subclasses separately (Sst, Pvalb, Vip, Lamp5) (Within subclass Patch-seq RNA preprocessing section). Finally, to measure prediction performance using the expression levels of the ion channels alone, we used only the ion channels expression from the 3,220 interneurons with subclass effects regressed out and constructed statistical models to predict ephys response.

### Prediction of electrophysiological features from gene expression patterns (across subclass)

For our across subclass prediction experiments, we iteratively selected one of the four mouse subclasses (e.g. Sst) as the training dataset, and combined the remaining three subclasses (e.g. Pvalb, Vip, Lamp5) to form a test dataset. To ensure the results were comparable to the all cells and within subclass experiments, we repeated the same cross validation strategy as the all cells and within subclass experiments to train multiple Sst models (across folds and seeds), but always evaluated on the entire test dataset (instead of the three outer test folds). Note that for the across subclass experiments, we use each subclass’s across subclass preprocessed data (**Cross subclass IPFX core and PCA feature preprocessing**).

### Identification of ephys response gene sets for heritability analysis

For each well-predicted ephys features in our within subclass experiments (Spearman rho > 0.4), we extracted the set of genes selected by the trained GLM to predict the ephys feature, termed the ephys (response) gene set. Because the within subclass performance is measured via cross validation, to extract the ephys genes we trained GLMs with alpha pre-set to 0.5 using the cv.glmnet function in the glmnet package^72^ in R. All genes with non-zero coefficients from the GLM were identified as the ephys genes. Because the IPFX PCA features consist of sets of ephys features (there are potentially multiple related PCs), we defined the ephys gene set as the union of the predictive genes for each of the PCs.

In our heritability calculations below, we computed permutation-based p-values by constructing up to 10,000 permuted gene sets per ephys gene set. Here we describe the procedure to generate one permutation, which we then repeat to generate multiple permutation gene sets. For each subclass, we permuted the identity of the neurons in the expression data but not the ephys data; this permutation scheme destroys the paired nature of the ephys and gene expression data, but maintains the gene-gene and ephys feature-feature correlation structure of the data. We used cross validation with alpha set to 0 to identify the lambda with largest cross validation accuracy, then trained a GLM with alpha set to 0 at the identified lambda value, and ranked the genes by coefficient magnitude. Among the selected genes, we mapped them to their 1-1 human ortholog, and discarded those genes without chromosome locations in GRCh37. Then, for each IPFX core and PCA feature, we selected the top K genes to form the permuted version of that IPFX feature’s ephys gene set, where *K* is set to the same size as the true IPFX feature’s ephys gene set. In the specific case of the IPFX PCA features, we extracted the same number of genes as the size of ephys response gene sets for the individual PCs, then took the union of the genes selected for each individual PC to form one permutation set for each IPFX PCA feature. We adaptively constructed up to 10,000 permuted gene sets per IPFX feature, such that more permuted gene sets were constructed if the permuted p-value was small.

### Partitioned heritability estimation procedure

We performed heritability analysis of multiple psychiatric and mental disorders using several different genomic annotations, including open chromatin regions for different interneuron classes^60^ (**Supplementary Table 4**), and regions associated with our ephys response gene sets. The procedure for estimating heritability and the associated significance is the same; the only difference in the calculations is in construction of the genome annotations. Here we describe our procedure for estimating heritability, assuming a genome annotation has been provided. Details of how we construct the annotations are found in later sections.

We first estimated the LD scores using a single genome annotation (details below), using a modified version of the baseline annotations (version 2.2) (https://alkesgroup.broadinstitute.org/LDSCORE) with the recommended 1cM window size. More specifically, we removed two annotations from the baseline annotations, MAF_Adj_Predicted_Allele_Age and MAF_Adj_LLD_AFR, due to the strong significant negative enrichments that we observed in the LD score partitioned heritability when using the Roadmap H3K27ac peaks. We then estimated the partitioned heritability from the generated LD scores using European 1000 Genome Phase 3 genotypes, frequency and LD score regression weight files downloaded from https://alkesgroup.broadinstitute.org/LDSCORE/ under the name: 1000G_Phase3_plinkfiles.tgz, 1000G_Phase3_frq.tgz, and 1000G_Phase3_weights_hm3_no_MHC.tgz respectively.

### Generation of ephys response gene set annotations for heritability analysis

We constructed genome annotation files for each ephys feature by applying the make_annot script to our ephys response gene sets’ chromosomal location with the recommended settings (window size = 100kb) and 1000 Genomes phase 3 genotype files. We also performed partitioned heritability on all of the permutation gene sets for each of the ephys features using the same procedure above. For each of the features, we count the number of permutation gene sets that have a smaller heritability enrichment p-value than the nominal heritability enrichment p-value attained by the true ephys gene set, and divide by the number of permutation sets. This number is then reported as the permutation-based p-value.

### Generation of subclass-specific open region annotations for heritability analysis

scATAC-seq peaks^60^ called from isocortex-derived SST, PVALB, LAMP5 and VIP neurons were downloaded from Supplementary Data 7 of https://www.nature.com/articles/s41588-020-00721-x#Sec47 in November 2021. The peaks were filtered for FRD significance (< 0.05) for each neuronal subclass and converted into BED format and subsequently mapped to hg37 coordinates using the UCSC liftover tool^73^ (https://genome.ucsc.edu/cgi-bin/hgLiftOver). The preprocessed peaks for each subclass were then compared with other subclasses to quantify the subclass-specificity of each peak by computing its frequency of occurrences across subclasses. Then based on each peak’s frequency of occurrences across subclass and which of the four subclasses the peak is enriched in, we assign peaks to subclass and generated multiple open chromatin annotations using the make_annot script for each of the four subclasses using peaks with frequency of occurrences between 1-4 subclasses.

### Gene co-expression network construction

For each IPFX core and PCA ephys feature, we constructed a gene co-expression network by calculating a pairwise gene-gene Pearson correlation matrix using the batch effect-corrected Patch-seq gene expression matrix, subsetted to the Sst subclass. We then binarized the correlation matrix using a 99.99^th^ percentile threshold, followed by subsetting of the genes to the union of the ephys feature-specific ephys gene set. For our visualization of the first AP dV/dt genes together with the ion channels (**Fig. 4a**), we included the set of all annotated ion channel genes during the subset operation to the first AP dV/dt ephys genes.

### Gene Ontology enrichment

For all of the 42 features shown in **Figure 1a**, we extracted the genes that are co-expressed with other genes (connected genes) from the binarized the correlation matrix. For each of the features, we then used the set of connected genes to performed gene ontology enrichment analysis with the C5 ontology annotations from MSigDb collections (version 7.5.1)^74^ using msigdbr and cluster profiler^75^ R packages while conditioning on the entire set of ephys genes for that feature as a background gene set. P-values from the result are corrected using FDR correction. We repeated this procedure of selecting connected genes from the co-expression network of the ephys gene set for each ephys feature, and performing gene ontology enrichment analysis with the C5 ontology annotations from MSigDb collections (version 7.5.1)^74^ using msigdbr and cluster profiler^75^ R packages while conditioning on the entire ephys response gene set as the background gene set. P-values from the result are corrected using the Benjamini-Hochberg procedure.

### Differential gene expression computation

From the mouse Patch-seq QCed count matrix, we identified all genes expressed in more than 10% of the neurons. We used the edgeR package version 3.26.8 in R with a glm model^76^ with default parameters to calculate differential expression, and included age, biological sex, hemisphere, cell type effects in the glm model.

### Kirrel3 coexpression network construction

After following the Within subclass and across subclass Patch-seq RNA preprocessing strategy for processing RNA data, we computed four subclass-specific, genome-wide gene-gene correlation matrices. For the Lamp5 and Pvalb correlation matrices, we first set the diagonal of the correlation matrix to zero, followed by binarization of the correlation matrix using the 99.95^th^ percentile as a threshold. Next, for each subclass separately, we subsetted the binarized gene gene correlation matrix using the top 100 correlated genes (correlation ephys genes) of the Frequency vs current slope and AP dV/dt PC1 ephys features. Finally, we visualized the gene co-expression network from the subsetted correlation matrix using the igraph package^77^ in R. For **Figures 5g-h**, we retained only nodes that are co-expressing with Kirrel3 gene in the Lamp5 and Pvalb co-expression networks.

### Kirrel3 gene expression and ephys correlation

After following the Within subclass and across subclass Patch-seq RNA and ephys preprocessing strategy, we extracted Kirrel3’s expression values, AP dV/dt PC1, and Frequency vs current slope values for both Pvalb neurons and Lamp5 neurons. We then plotted a scatter plot between Kirrel3 expression and the two ephys feature values for both Lamp5 and Pvalb cell types (**Fig 5 a-d**). We further fitted a linear model on scatter plot and noted the F-test P-value on the fitted linear model.

### Adhesion molecules - ephys correlation pattern analysis

We retrieved a list of cell-cell adhesion genes (GO: 0098609) from http://www.informatics.jax.org/go/term/GO:0098609 on June 2022. Using the RNA and ephys data from Within subclass and across subclass Patch-seq RNA and ephys preprocessing, we computed the gene – ephys Pearson correlations for the 42 ephys features shown in **Figure 1** for each of the four subclasses. We then retained the top 100 genes that are correlated to each of the 42 ephys features (correlation-ephys genes) for each of the four subclasses, and further subsetted to adhesion molecules that belong in GO: 0098609 that were expressed in more than 10% of the neurons in the mouse Patch-seq QCed count matrix. For each of the adhesion molecules that passed the 10% expression threshold, we checked if it is a correlation-ephys genes for each of the 42 ephys features across four cell types, and retained only the adhesion genes that were correlation-ephys genes for more than 6 features (high confidence adhesion molecules).

### Function switching inference of cell adhesion genes

To infer function switching of the individual cell adhesion genes, we first assigned the 42 ephys features into six function categories based on **Supplementary Table 1**. For high confidence adhesion molecules, we counted the number of ephys features within each of the function categories that the gene is a correlation-ephys gene of. Each adhesion molecule is then assigned one or more dominant function categories in each subclass, based on which categories the adhesion molecule has more than 3 correlated ephys features with. An adhesion molecule is defined to exhibit function category switching if the dominant function categories in any subclass differs from the dominant function categories in another subclass, given both subclasses have at least one dominant category. Furthermore, if the switching behavior involving two subclasses is such that the dominant categories are mutually exclusive between the two subclasses, we classify the adhesion gene as a binary switching gene.

---

**Fig. S1.**
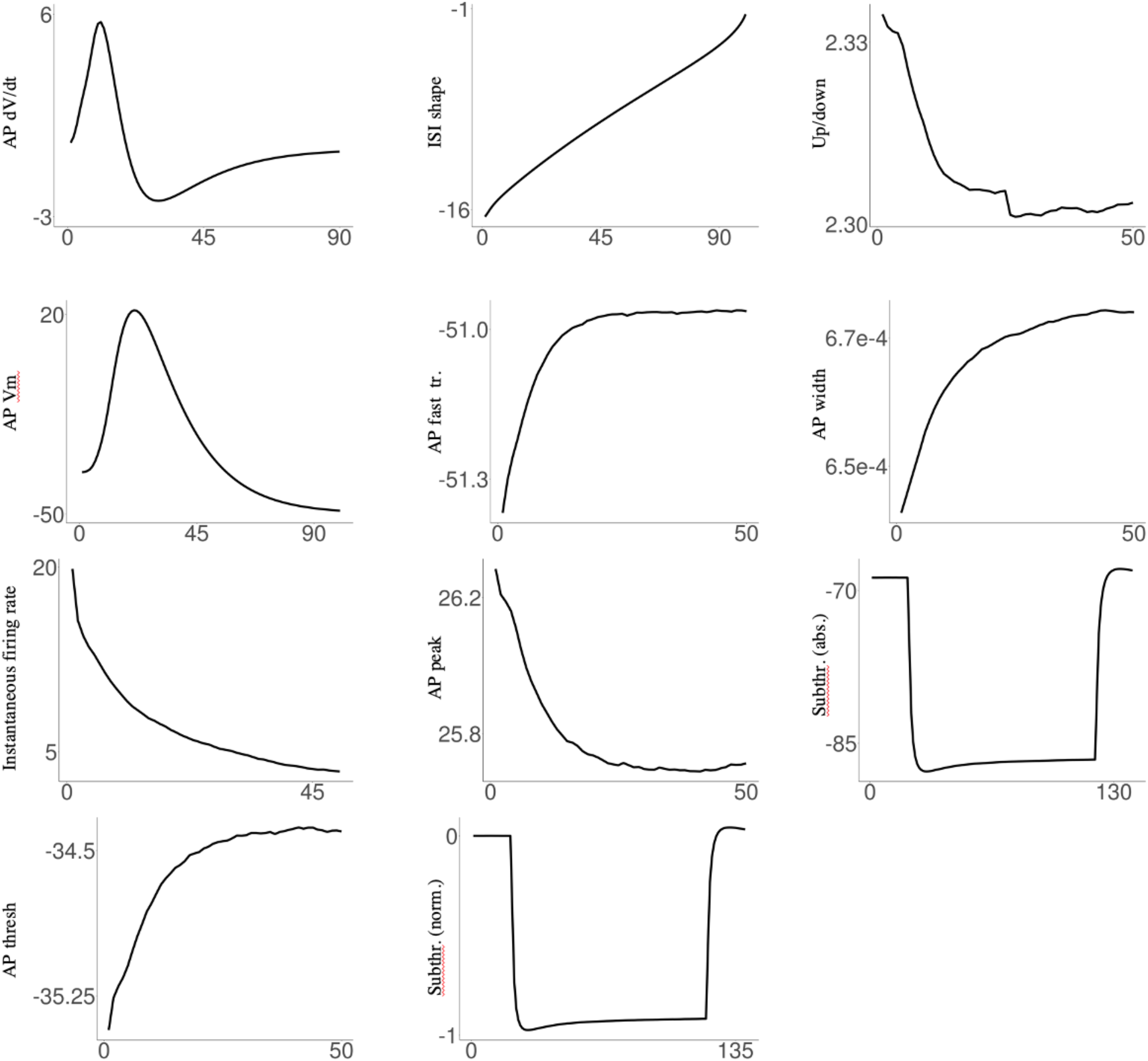
Visualization of the 11 IPFX PCA feature vectors. Plot illustrate the average feature vectors used to calculate the IPFX PCA ephys features for Sst, Pvalb, Lamp5 and Vip. The x-axis represents time. For a subset of features (AP Width, Up/down, AP fast tr., Inst. firing rate, AP thresh., AP peak), only the ephys response from rheobase stimulation is plotted. For a subset of features (Subthr. (norm) and Subthr. (abs)), only the ephys response from the first hyperpolarizing current step is shown.

**Fig. S2.**
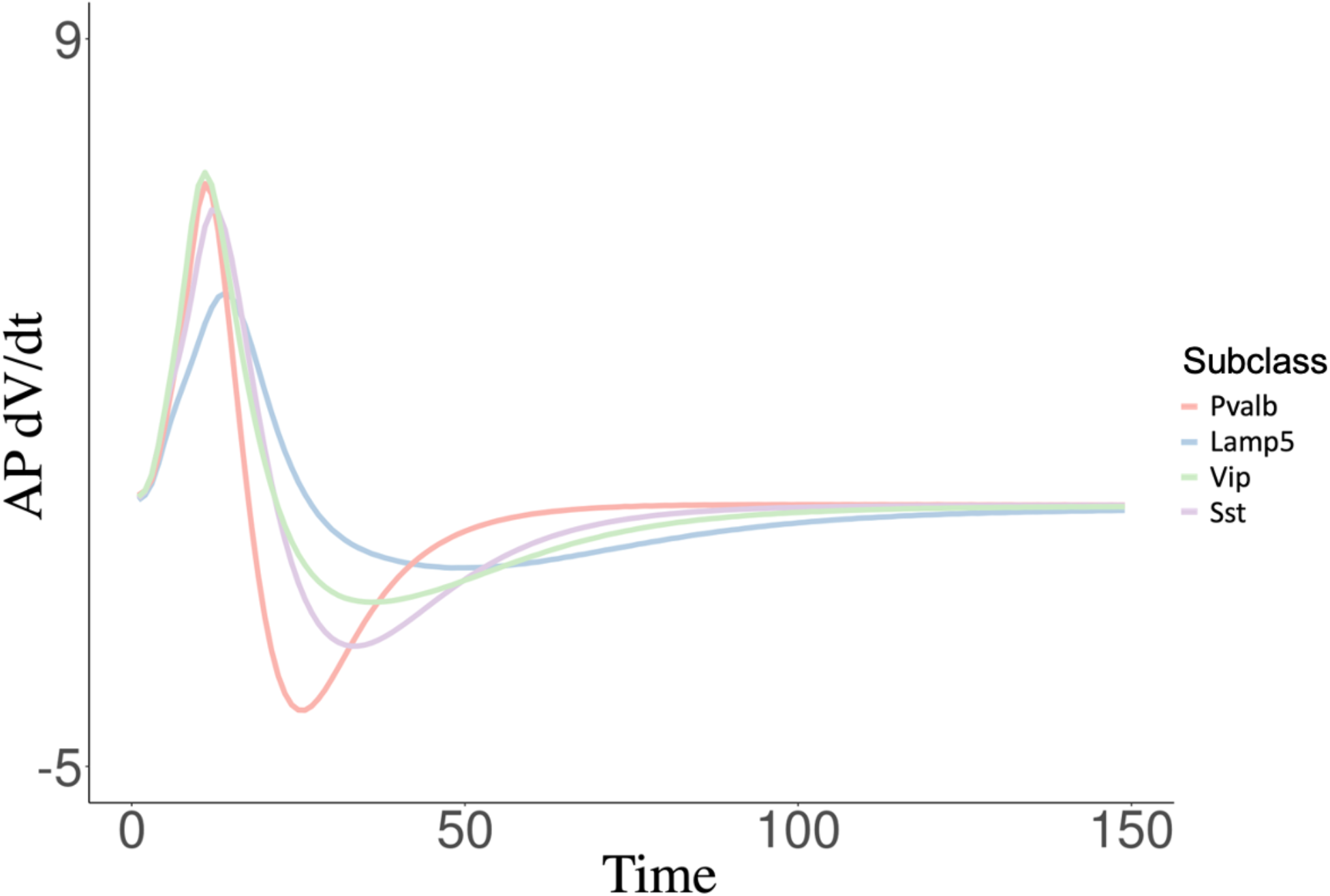
Average AP dV/dt measurements over time for each of the four interneuron subclasses. Values are averaged across all neurons of the same subclass.

**Fig. S3.**
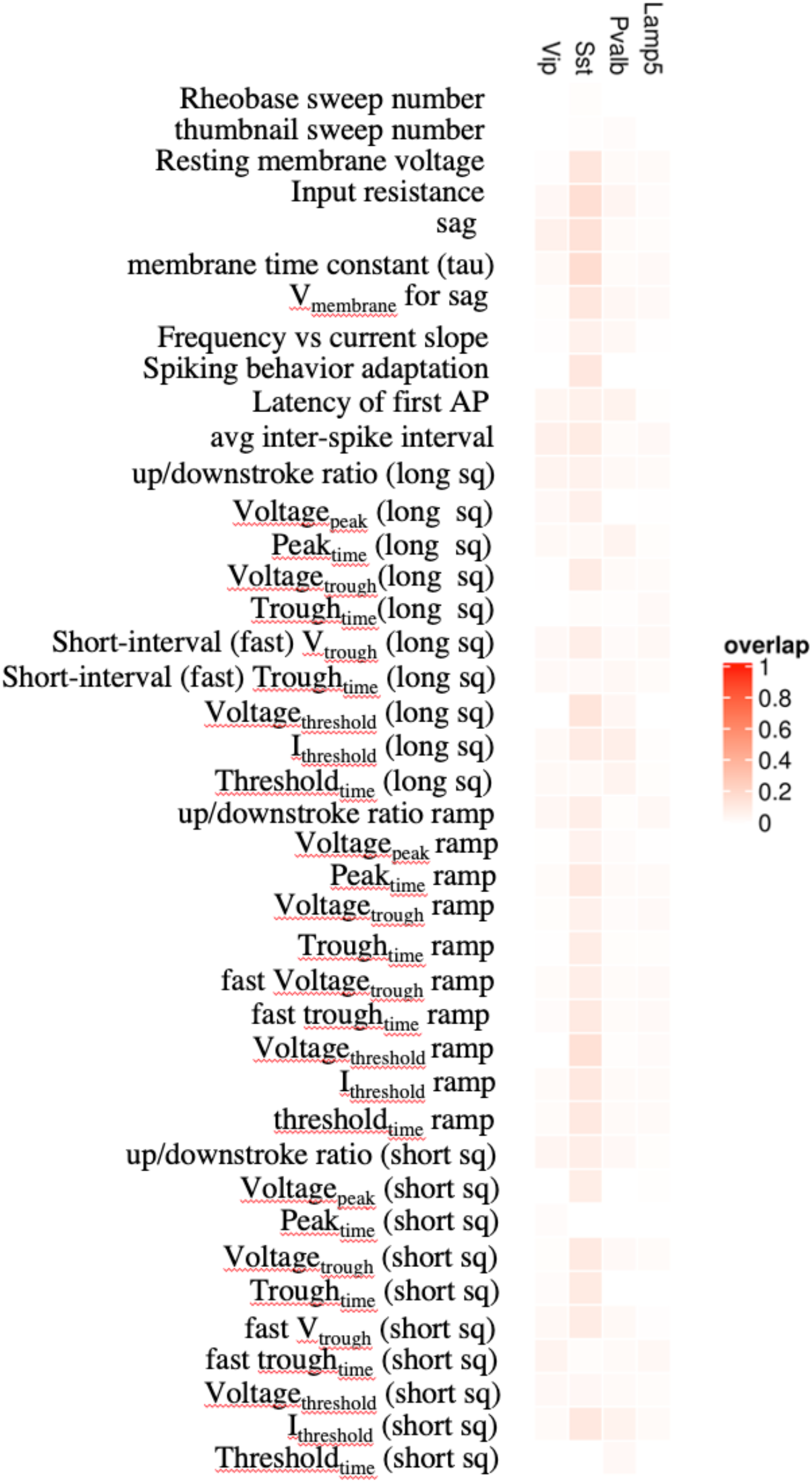
Overlap of ephys genes between subclass-specific models and all cells model. Heatmap represents the Jaccard index between the ephys genes identified by the subclass-specific models and the all cells model across all IPFX core ephys features.

**Fig. S4.**
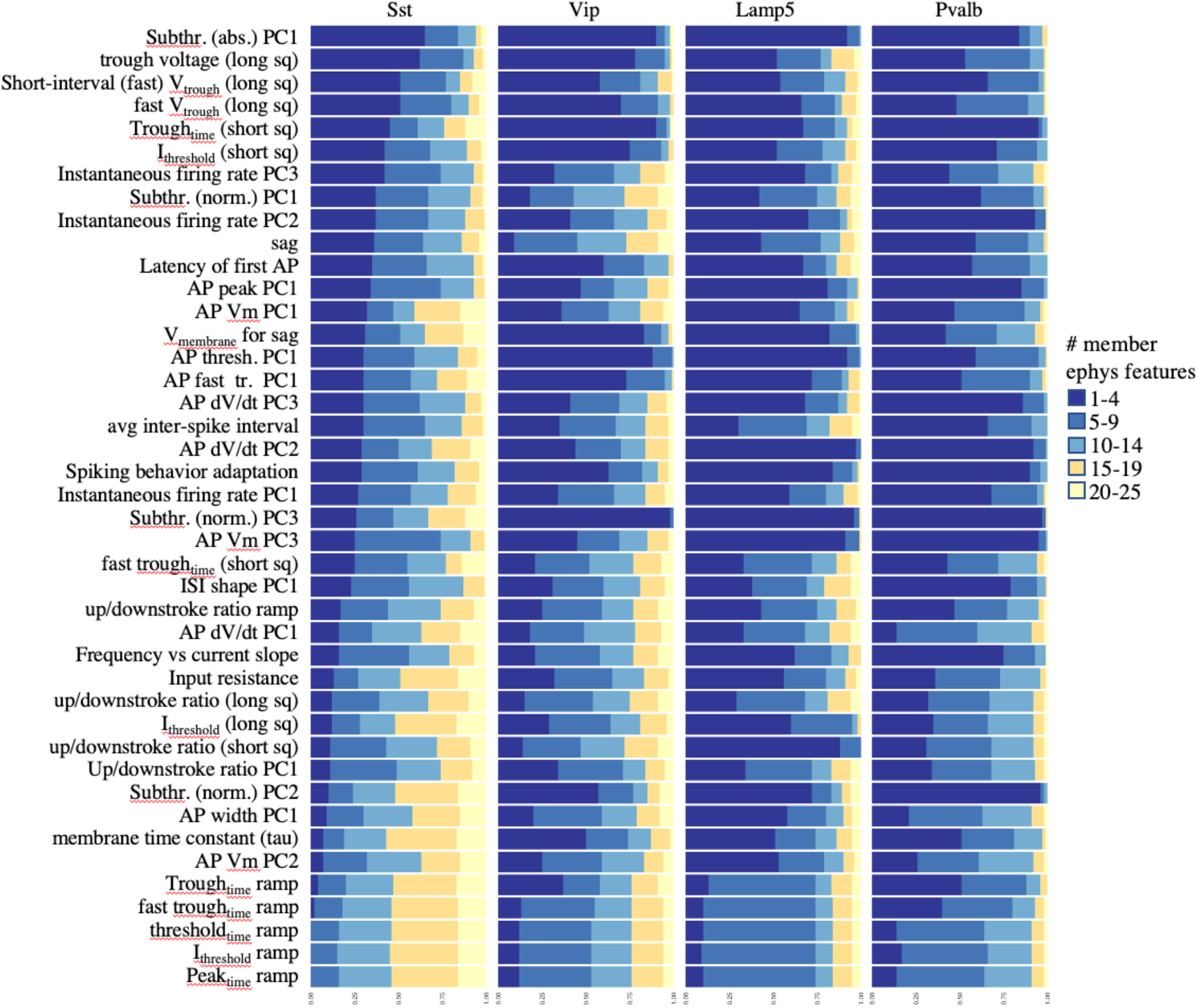
Ephys response genes are ephys feature-specific in each of the four subclasses. Stacked bar charts indicating, for each ephys feature in each of the four subclasses, the fraction of the corresponding correlation-ephys genes that are also correlation-ephys genes for other ephys features in Sst. Correlation-ephys genes are grouped based on how many other ephys features they are correlated with. Features are sorted based on average number of other ephys features that each ephys gene is correlated with in the SST subclass.

**Fig. S5.**
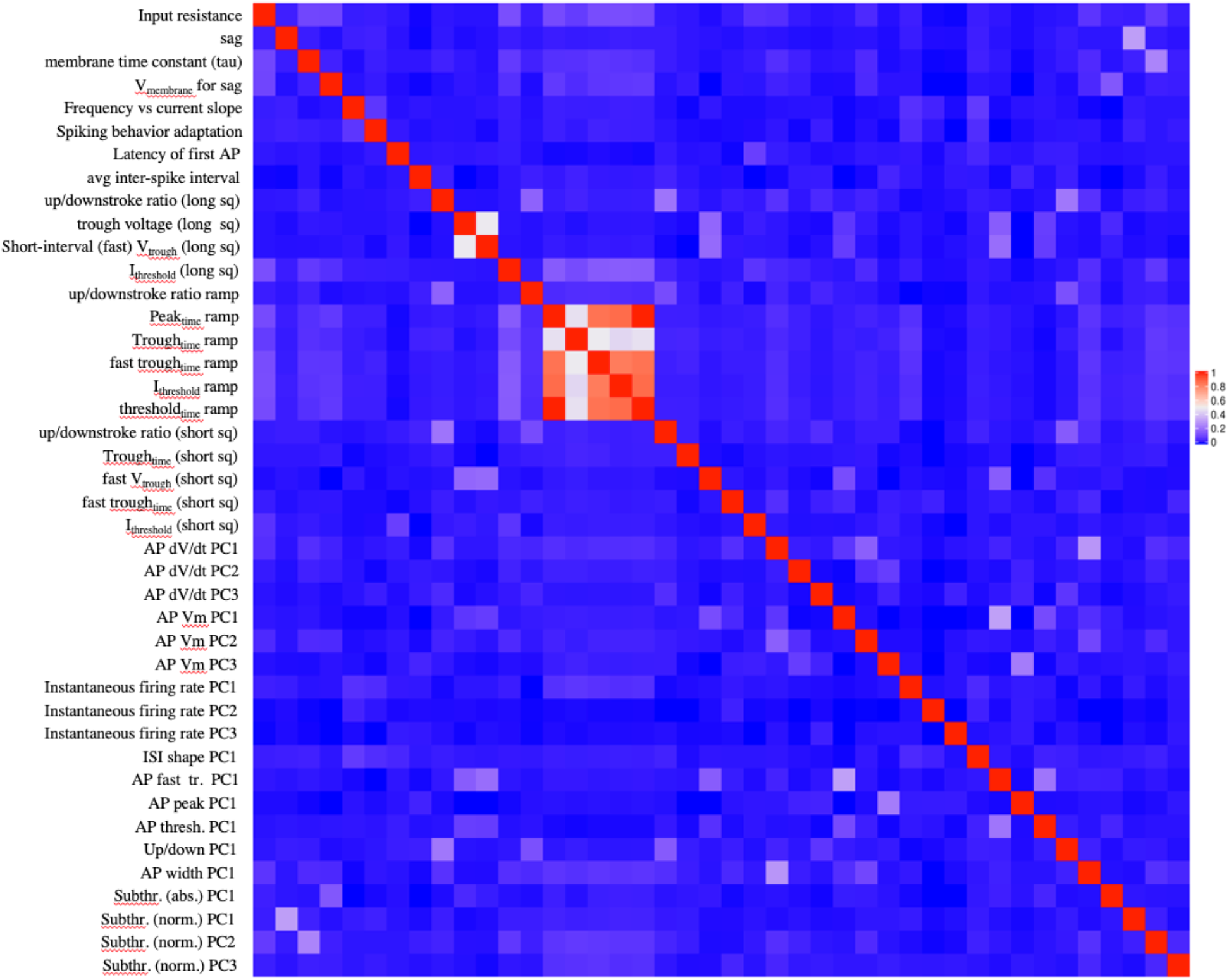
Overlap of ephys genes identified between ephys features of Sst. Overlap is calculated as the Jaccard index between ephys gene sets of pairs of ephys features, where the ephys gene sets are identified as all genes whos GLM coefficients are non-zero, where the GLM is trained with alpha = 0.5.

**Fig. S6.**
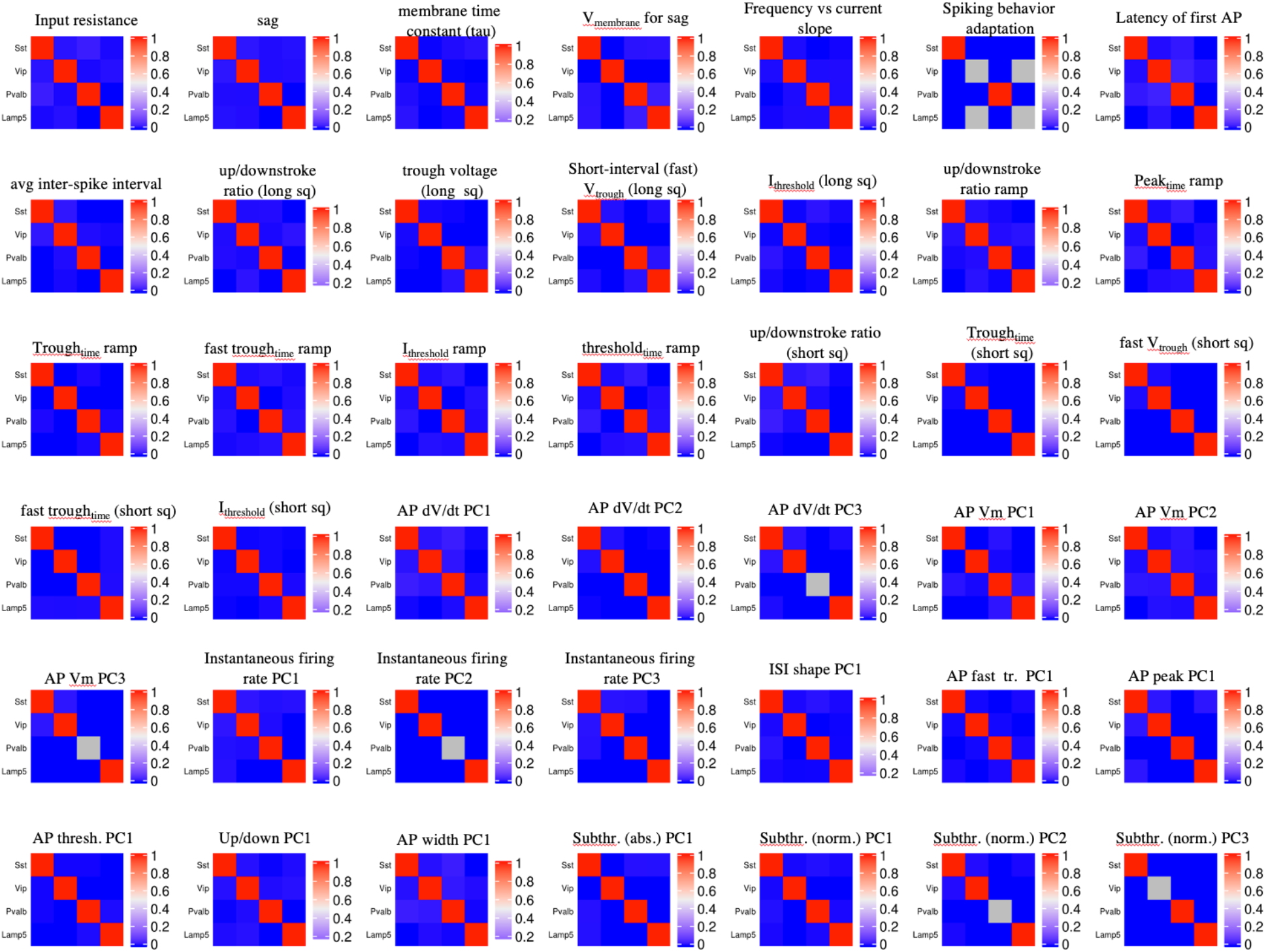
Overlap of ephys genes identified between different subclasses. Each heatmap indicates the Jaccard index of ephys gene sets identified for the same ephys feature but different subclasses.

**Fig. S7.**
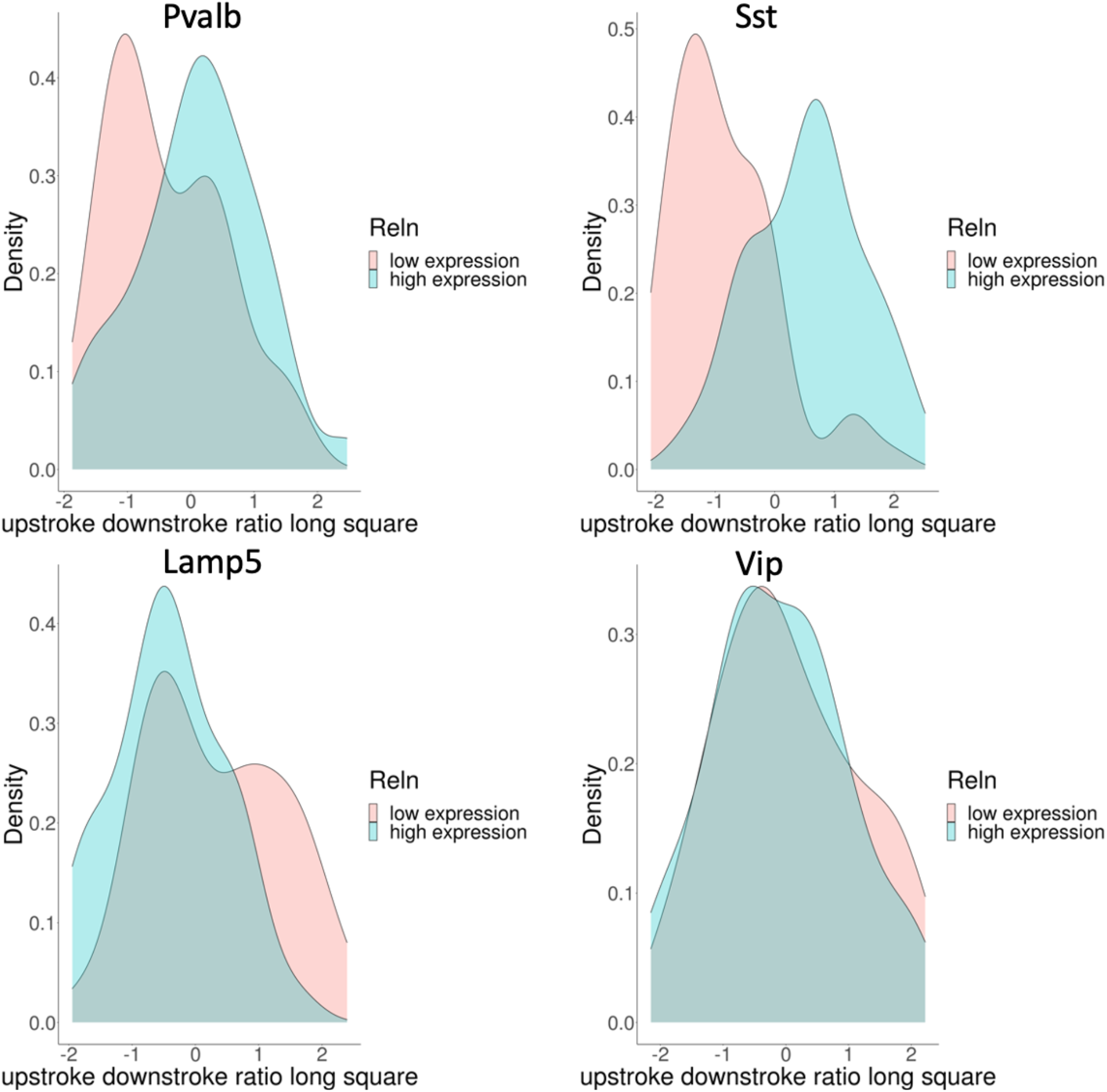
Density plots of the Up/downstroke ratio (long sq) ephys feature for the four subclasses, as a function of Reln expression. For each subclass, the up/downstroke ratio value distributions are divided into two distributions based on the relative expression of Reln in the same neuron. Reln low expression refers to neurons in which Reln expression is less than the 10^th^ percentile of Reln expression (low expression), and high expression refers to neurons in which Reln expression is greater than the 90^th^ percentile of Reln expression within each of the four subclasses.

**Fig. S8.**
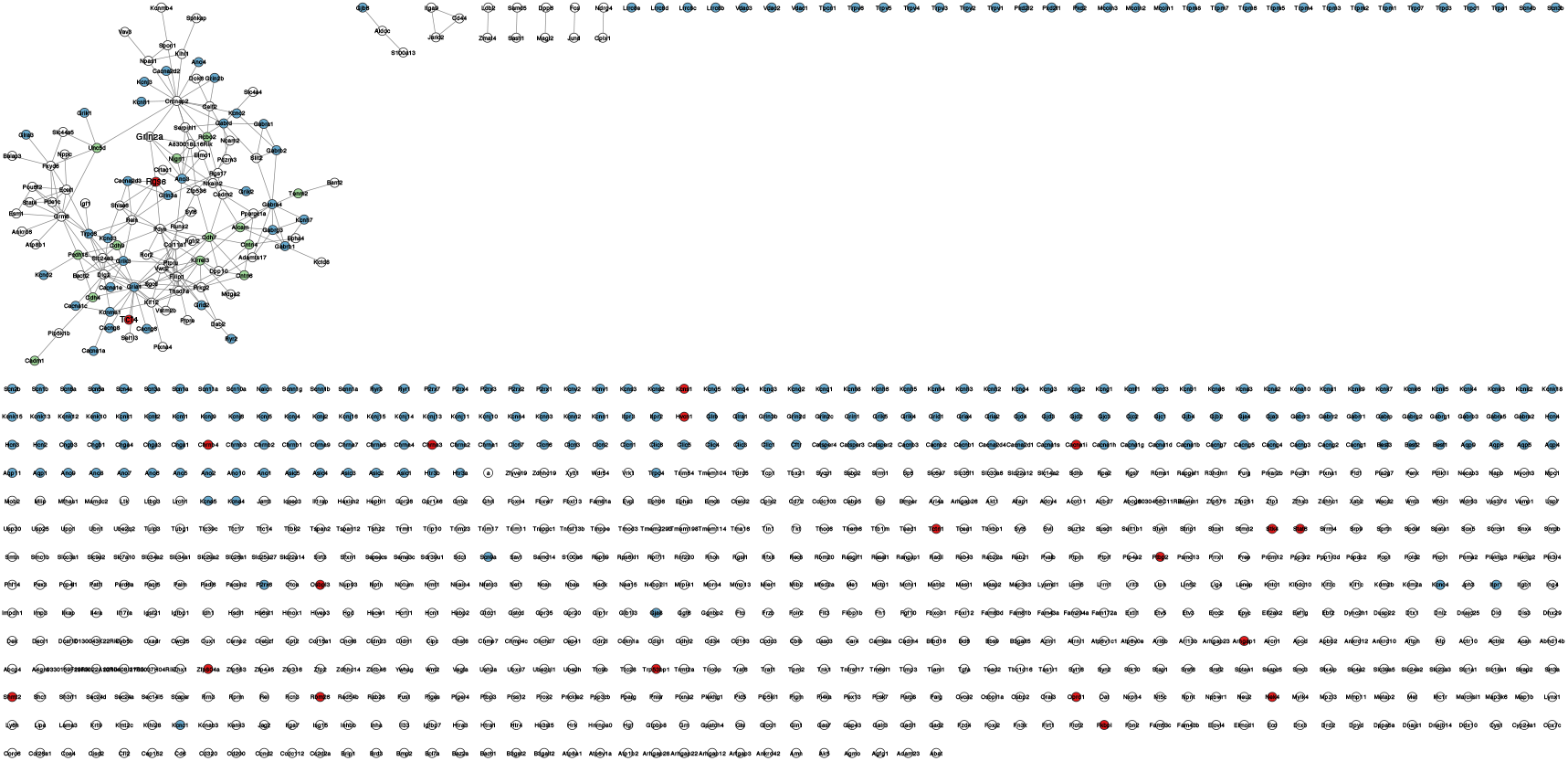
Full co-expression network of Sst AP dV/dt genes and ion channels. Co-exprsesion etwork showing gene-gene correlations for the union of genes within the Sst AP dV/dt gene set and ion channels. Edges represents genes with relatively high correlation (>99.99^th^ percentile). Red nodes represent genes with genome wide significant SNPs in psychiatric GWAS (**see Methods**) within 5kb of the gene TSS. Blue nodes represent ion channels; green nodes represent adhesion molecules in annotated within GO:0098742.

**Fig. S9.**
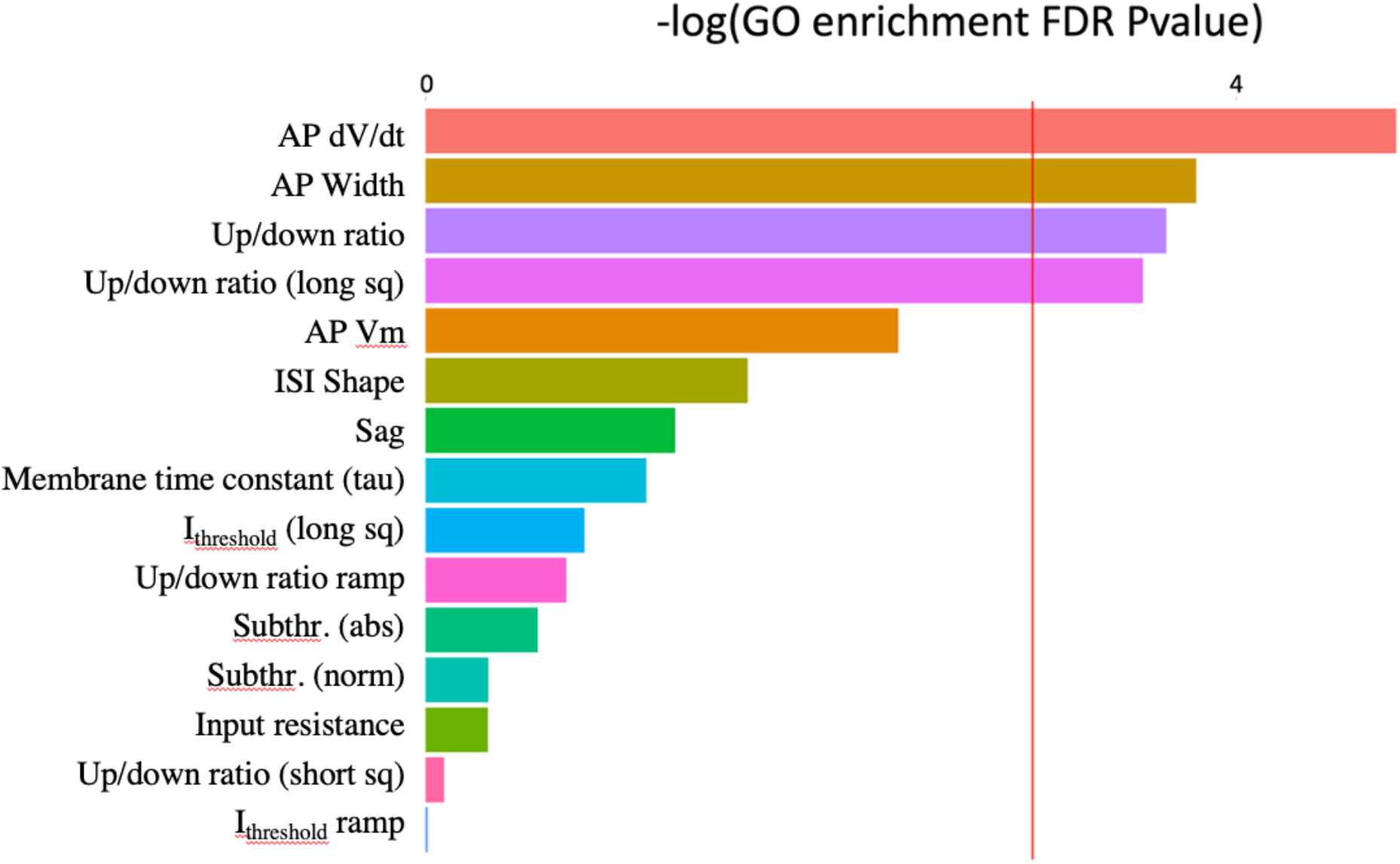
Gene ontology enrichment of ephys gene sets for selected Sst ephys features. Bar chart indicates the negative log of FDR corrected P-values for enrichment of ephys genes with respect to GO:0098742 (cell-cell adhesion via plasma-membrane adhesion molecules). Ephys features are restricted to those used for heritability analysis. Red horizontal line represents significance (FDR corrected P-value = 0.05).

**Fig. S10.**
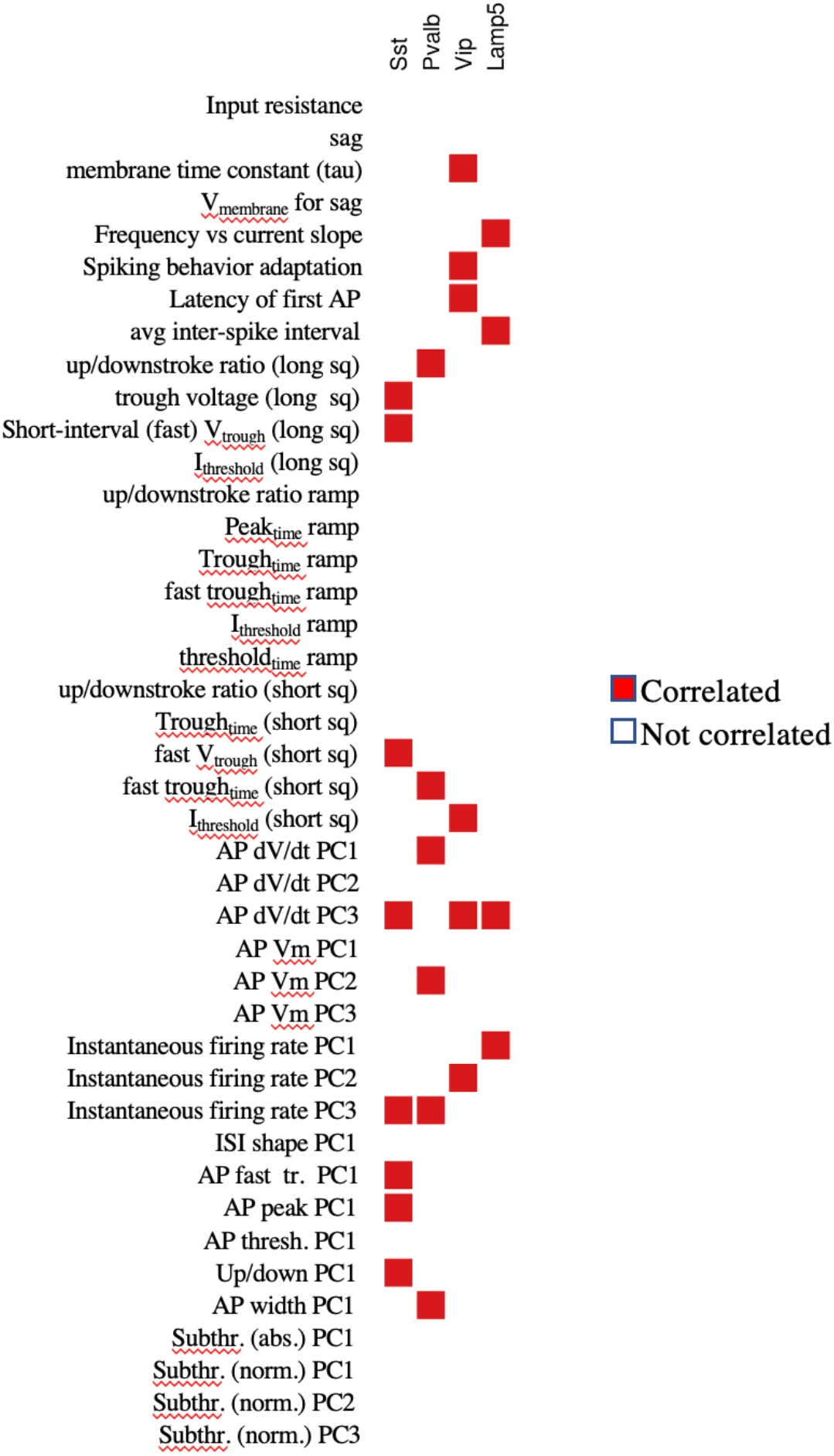
Ephys associations between Kirrel3 and different ephys features across the four subclasses. A red square indicates that Kirrel3 was a correlation-ephys gene (top 100 most correlated genes) for a given ephys feature in a given subclass.

**Fig. S11.**
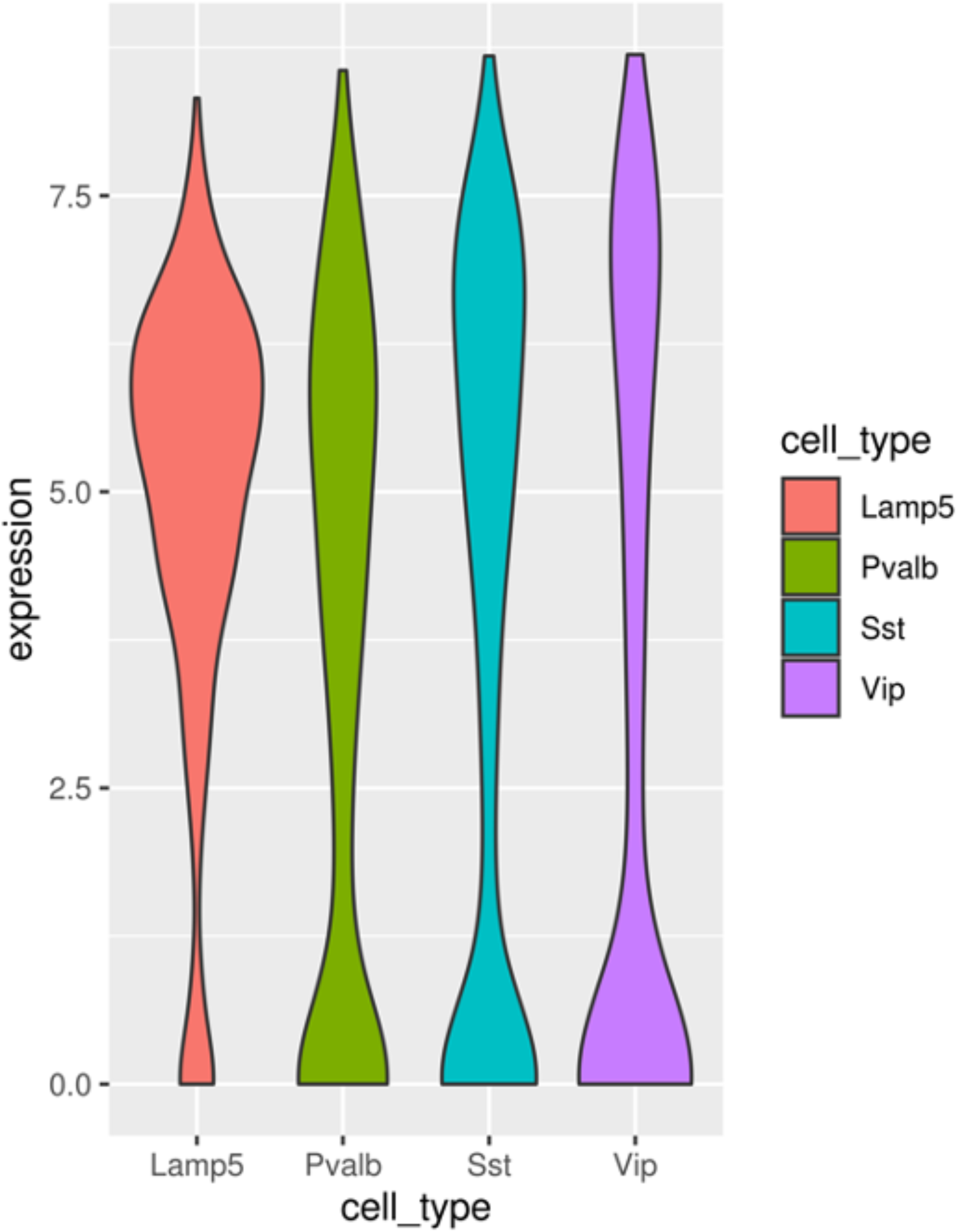
TPM expression levels of Kirrel3 in the four subclasses after QC.

**Fig. S12.**
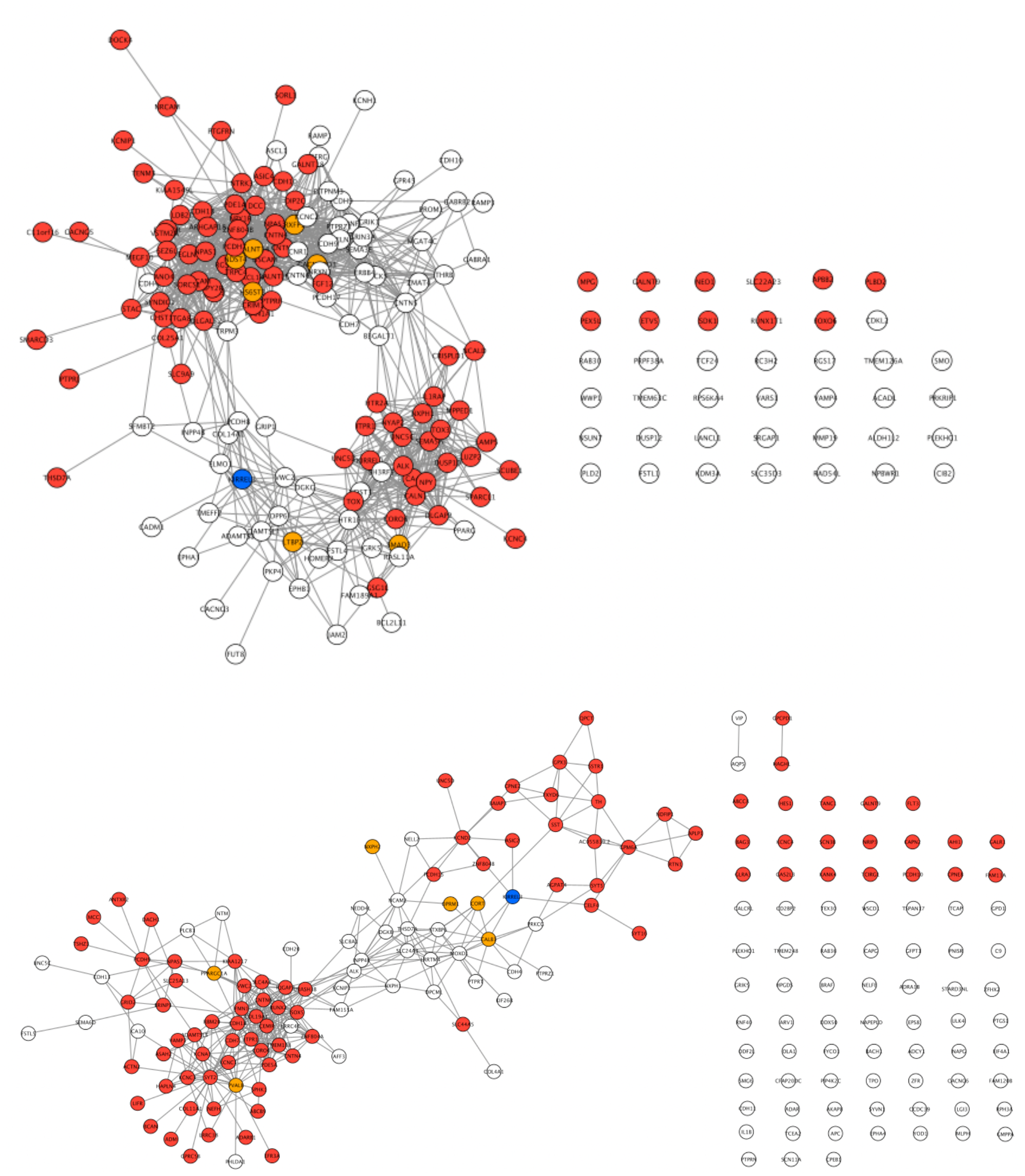
The entire co-expression network of Kirrel3 shown in Figure 5g, h. Each co-expression network includes the correlation-ephys genes (top 100 most correlated) for the Frequency vs current slope (white node), AP dV/dt PC1 (red node) and both (orange node) ephys features for Lamp5 (top) and Pvalb (bottom). Edges are binarized using a 99.95^th^ percentile cutoff of the genome-wide gene-gene correlation matrix.

**Fig. S13.**
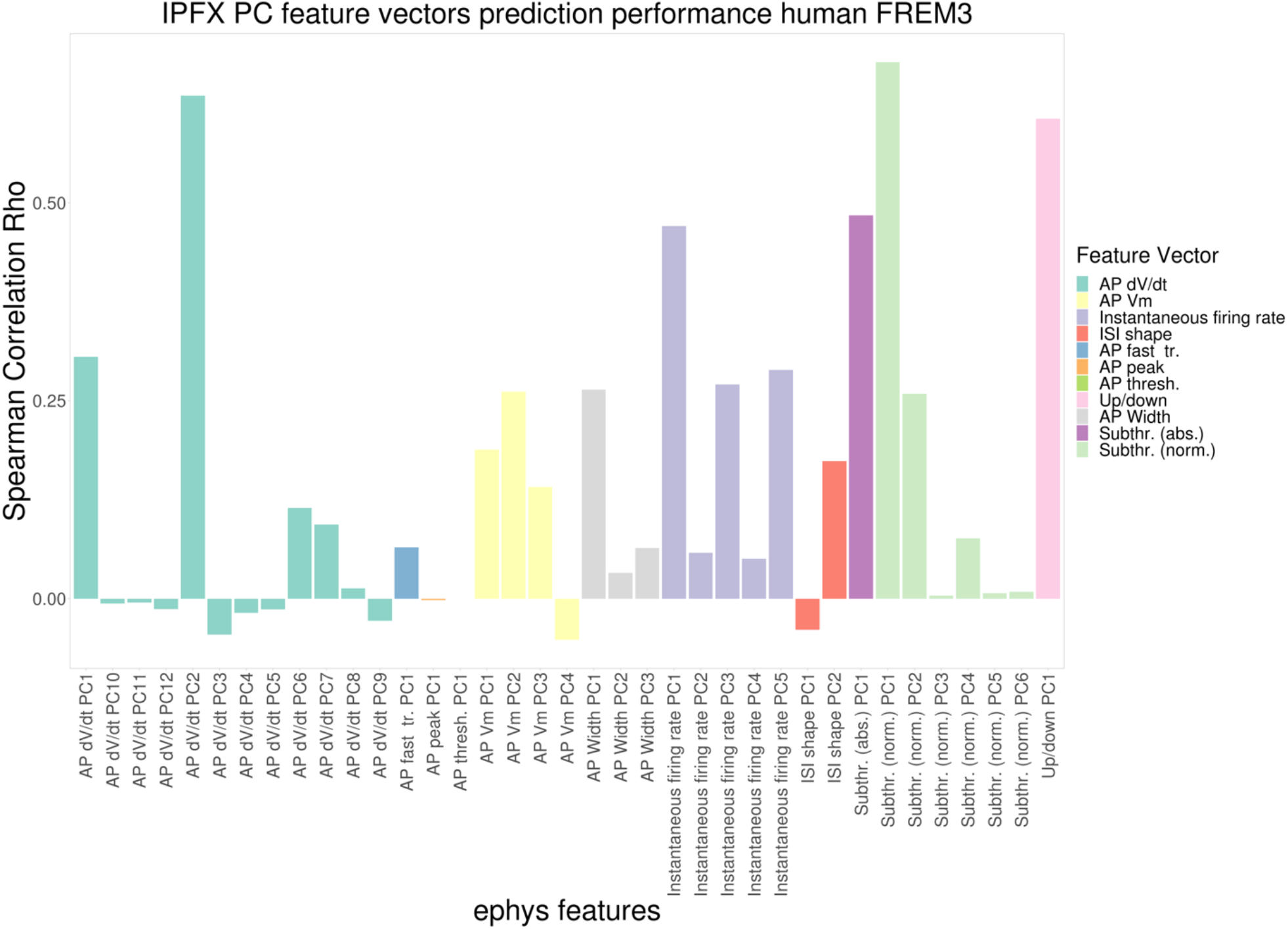
Cross-validation performance of predictive models trained on human FREM3 neurons, and evaluated on IPFX PCA ephys features. Each bar represents a different IPFX PCA ephys feature measured on the FREM3 neurons. Bars are colored by feature vector.

**Fig. S14.**
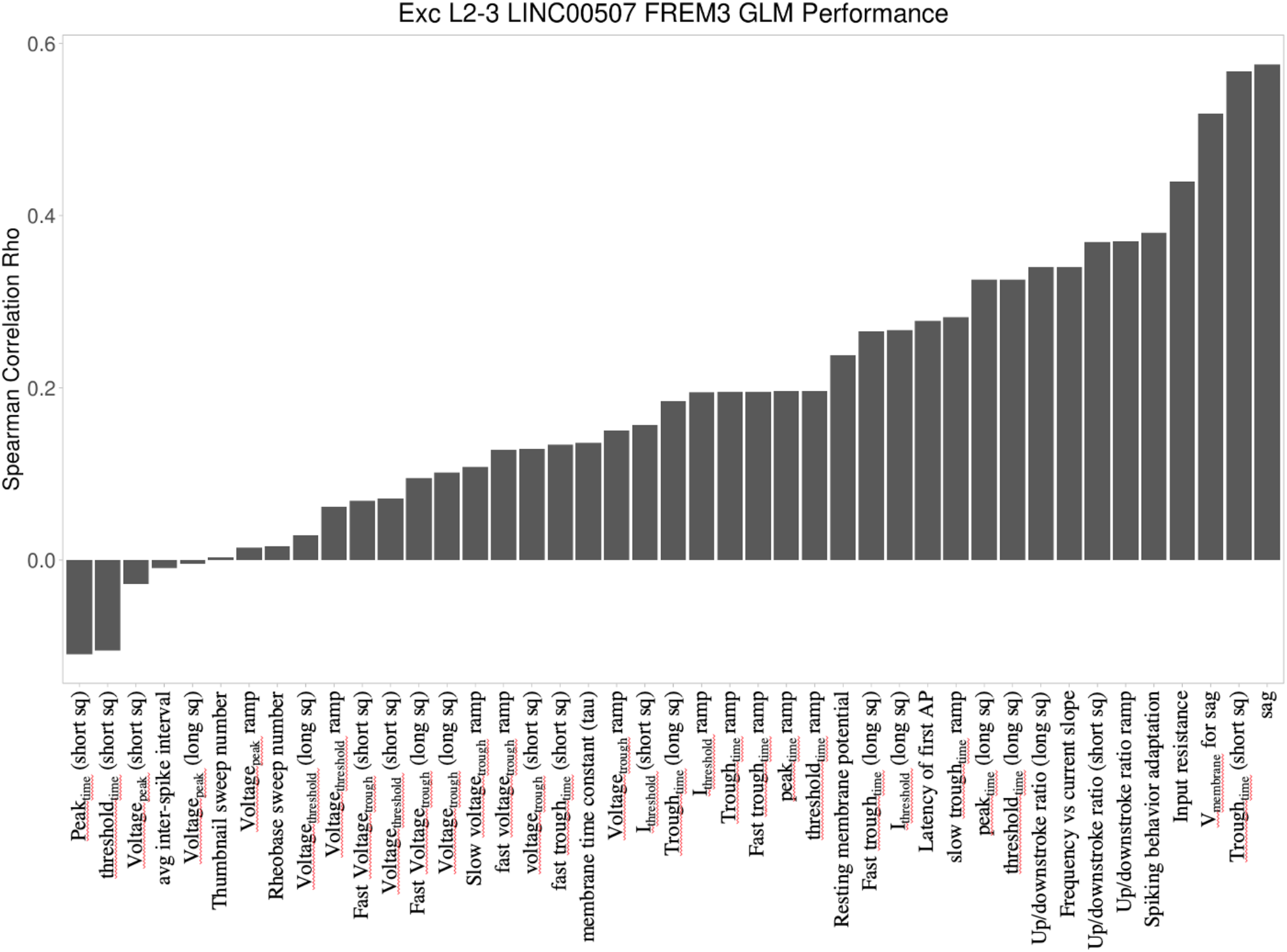
Cross-validation performance of predictive models trained on human FREM3 neurons, and evaluated on IPFX core ephys features. Each bar represents a different IPFX core ephys feature measured on the FREM3 neurons.

**Fig. S15.**
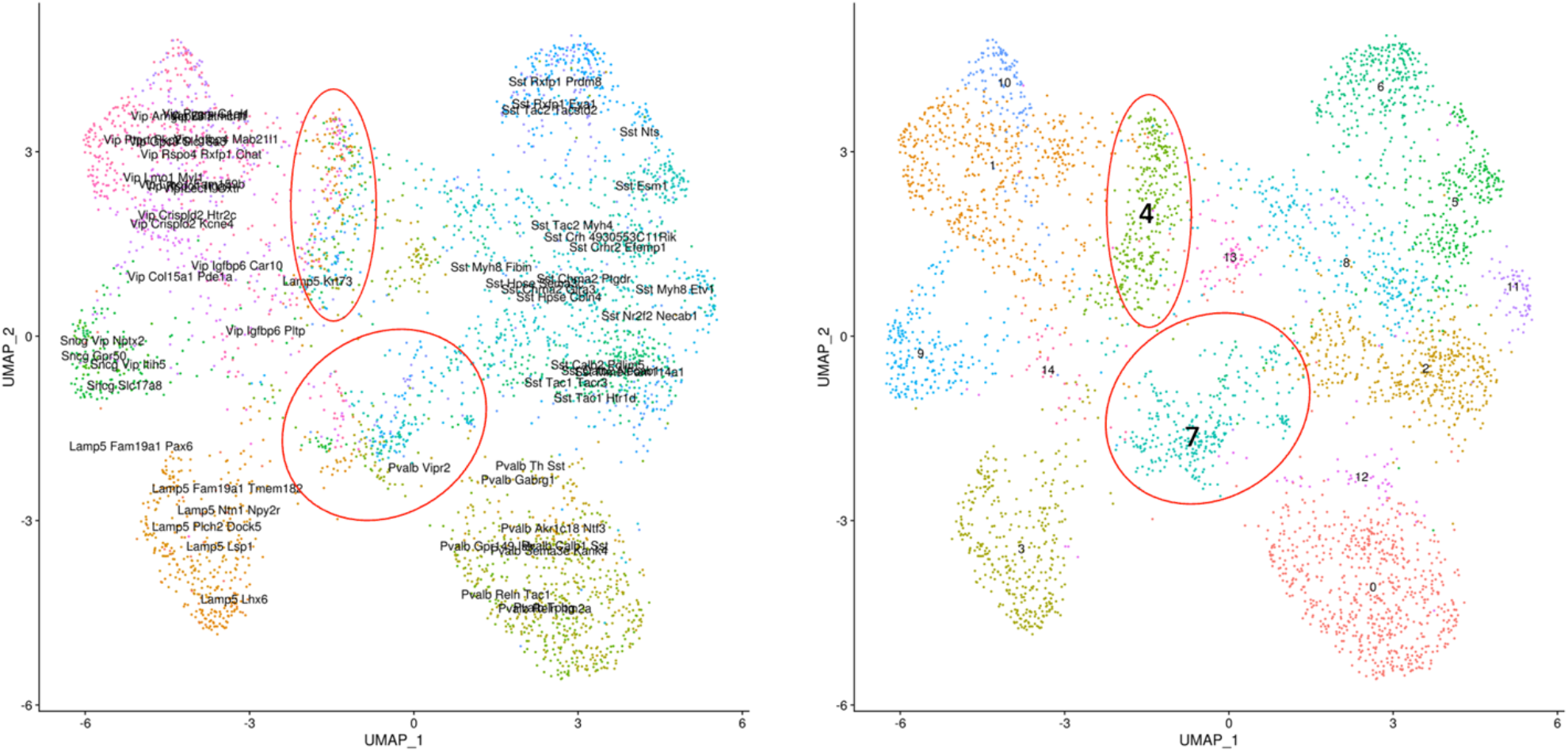
UMAP visualization of Patch-seq RNA samples. UMAP plot visualizes the mouse Patch-seq expression profiles colored by cell types (left) and cluster number (right). The UMAP is generated using the top 40 PCs of the expression data. Cell cluster labels are generated using the shared nearest neighbor implemented in the Seurat package. Cluster 4 and 7 (circled in red) show heterogeneous mixture of cell types and were subsequently removed for quality control.

## REFERENCES

1. Consortium, T. S. W. G. of the P. G., Ripke, S., Walters, J. T. & O’Donovan, M. C. Mapping genomic loci prioritises genes and implicates synaptic biology in schizophrenia. 2020.09.12.20192922 Preprint at https://doi.org/10.1101/2020.09.12.20192922 (2020).

2. Ripke, S. et al. Biological insights from 108 schizophrenia-associated genetic loci. Nature 511, 421–427 (2014).

3. Ruderfer, D. M. et al. Genomic Dissection of Bipolar Disorder and Schizophrenia, Including 28 Subphenotypes. Cell 173, 1705-1715.e16 (2018).

4. Mullins, N. et al. Genome-wide association study of more than 40,000 bipolar disorder cases provides new insights into the underlying biology. Nat. Genet. 53, 817–829 (2021).

5. Stahl, E. A. et al. Genome-wide association study identifies 30 loci associated with bipolar disorder. Nat. Genet. 51, 793–803 (2019).

6. Howard, D. M. et al. Genome-wide meta-analysis of depression identifies 102 independent variants and highlights the importance of the prefrontal brain regions. Nat. Neurosci. 22, 343–352 (2019).

7. Nagel, M. et al. Meta-analysis of genome-wide association studies for neuroticism in 449,484 individuals identifies novel genetic loci and pathways. Nat. Genet. 50, 920–927 (2018).

8. Wray, N. R. et al. Genome-wide association analyses identify 44 risk variants and refine the genetic architecture of major depression. Nat. Genet. 50, 668–681 (2018).

9. Grove, J. et al. Identification of common genetic risk variants for autism spectrum disorder. Nat. Genet. 51, 431–444 (2019).

10. Demontis, D. et al. Discovery of the first genome-wide significant risk loci for attention deficit/hyperactivity disorder. Nat. Genet. 51, 63–75 (2019).

11. Significant Locus and Metabolic Genetic Correlations Revealed in Genome-Wide Association Study of Anorexia Nervosa | American Journal of Psychiatry. https://ajp.psychiatryonline.org/doi/full/10.1176/appi.ajp.2017.16121402.

12. Arnold, P. D. et al. Revealing the complex genetic architecture of obsessive–compulsive disorder using meta-analysis. Mol. Psychiatry 23, 1181–1188 (2018).

13. Nievergelt, C. M. et al. International meta-analysis of PTSD genome-wide association studies identifies sex- and ancestry-specific genetic risk loci. Nat. Commun. 10, 4558 (2019).

14. Yu, D. et al. Interrogating the Genetic Determinants of Tourette’s Syndrome and Other Tic Disorders Through Genome-Wide Association Studies. Am. J. Psychiatry 176, 217–227 (2019).

15. Sullivan, P. F. & Geschwind, D. H. Defining the Genetic, Genomic, Cellular, and Diagnostic Architectures of Psychiatric Disorders. Cell 177, 162–183 (2019).

16. Tansey, K. E. & Hill, M. J. Enrichment of schizophrenia heritability in both neuronal and glia cell regulatory elements. Transl. Psychiatry 8, 1–6 (2018).

17. Skene, N. G. et al. Genetic identification of brain cell types underlying schizophrenia. Nat. Genet. 50, 825–833 (2018).

18. Schijven, D. et al. Comprehensive pathway analyses of schizophrenia risk loci point to dysfunctional postsynaptic signaling. Schizophr. Res. 199, 195–202 (2018).

19. Gandal, M. J. et al. Transcriptome-wide isoform-level dysregulation in ASD, schizophrenia, and bipolar disorder. Science 362, eaat8127 (2018).

20. Walker, R. L. et al. Genetic Control of Expression and Splicing in Developing Human Brain Informs Disease Mechanisms. Cell 179, 750-771.e22 (2019).

21. Wockner, L. F. et al. Genome-wide DNA methylation analysis of human brain tissue from schizophrenia patients. Transl. Psychiatry 4, e339–e339 (2014).

22. Wang, D. et al. Comprehensive functional genomic resource and integrative model for the human brain. Science 362, eaat8464 (2018).

23. Gnanapavan, S. & Giovannoni, G. Neural cell adhesion molecules in brain plasticity and disease. Mult. Scler. Relat. Disord. 2, 13–20 (2013).

24. Hidese, S. et al. Cerebrospinal fluid neural cell adhesion molecule levels and their correlation with clinical variables in patients with schizophrenia, bipolar disorder, and major depressive disorder. Prog. Neuropsychopharmacol. Biol. Psychiatry 76, 12–18 (2017).

25. O’Dushlaine, C. et al. Molecular pathways involved in neuronal cell adhesion and membrane scaffolding contribute to schizophrenia and bipolar disorder susceptibility. Mol. Psychiatry 16, 286–292 (2011).

26. Triebelhorn, J. et al. Induced neural progenitor cells and iPS-neurons from major depressive disorder patients show altered bioenergetics and electrophysiological properties. 2021.04.30.441774 Preprint at https://doi.org/10.1101/2021.04.30.441774 (2021).

27. Ni, P. et al. iPSC-derived homogeneous populations of developing schizophrenia cortical interneurons have compromised mitochondrial function. Mol. Psychiatry 25, 2873–2888 (2020).

28. Page, S. C. et al. Electrophysiological measures from human iPSC-derived neurons are associated with schizophrenia clinical status and predict individual cognitive performance. Proc. Natl. Acad. Sci. 119, e2109395119 (2022).

29. Zhu, M. et al. Shank3-deficient thalamocortical neurons show HCN channelopathy and alterations in intrinsic electrical properties. J. Physiol. 596, 1259–1276 (2018).

30. Brandon, N. J. & Sawa, A. Linking neurodevelopmental and synaptic theories of mental illness through DISC1. Nat. Rev. Neurosci. 12, 707–722 (2011).

31. Gauthier, J. et al. De novo mutations in the gene encoding the synaptic scaffolding protein SHANK3 in patients ascertained for schizophrenia. Proc. Natl. Acad. Sci. 107, 7863–7868 (2010).

32. Brandon, N. J. et al. Understanding the Role of DISC1 in Psychiatric Disease and during Normal Development. J. Neurosci. 29, 12768–12775 (2009).

33. Zhou, Y. et al. Mice with Shank3 Mutations Associated with ASD and Schizophrenia Display Both Shared and Distinct Defects. Neuron 89, 147–162 (2016).

34. Nagy, C. et al. Single-nucleus transcriptomics of the prefrontal cortex in major depressive disorder implicates oligodendrocyte precursor cells and excitatory neurons. Nat. Neurosci. 23, 771–781 (2020).

35. Bryois, J. et al. Evaluation of chromatin accessibility in prefrontal cortex of individuals with schizophrenia. Nat. Commun. 9, 3121 (2018).

36. Hoffman, G. E. et al. CommonMind Consortium provides transcriptomic and epigenomic data for Schizophrenia and Bipolar Disorder. Sci. Data 6, 180 (2019).

37. Giusti-Rodríguez, P. et al. Using three-dimensional regulatory chromatin interactions from adult and fetal cortex to interpret genetic results for psychiatric disorders and cognitive traits. 406330 Preprint at https://doi.org/10.1101/406330 (2019).

38. Gusev, F. E. et al. Chromatin profiling of cortical neurons identifies individual epigenetic signatures in schizophrenia. Transl. Psychiatry 9, 1–10 (2019).

39. McKinney, C. E. Using induced pluripotent stem cells derived neurons to model brain diseases. Neural Regen. Res. 12, 1062–1067 (2017).

40. Hoffmann, A., Ziller, M. & Spengler, D. Progress in iPSC-Based Modeling of Psychiatric Disorders. Int. J. Mol. Sci. 20, 4896 (2019).

41. Wen, Z., Christian, K. M., Song, H. & Ming, G. Modeling psychiatric disorders with patient-derived iPSCs. Curr. Opin. Neurobiol. 36, 118–127 (2016).

42. Young-Pearse, T. L. & Morrow, E. M. Modeling developmental neuropsychiatric disorders with iPSC technology: challenges and opportunities. Curr. Opin. Neurobiol. 36, 66–73 (2016).

43. Brennand, K. J. et al. Modelling schizophrenia using human induced pluripotent stem cells. Nature 473, 221–225 (2011).

44. Volpato, V. et al. Reproducibility of Molecular Phenotypes after Long-Term Differentiation to Human iPSC-Derived Neurons: A Multi-Site Omics Study. Stem Cell Rep. 11, 897–911 (2018).

45. Li, L., Chao, J. & Shi, Y. Modeling neurological diseases using iPSC-derived neural cells. Cell Tissue Res. 371, 143–151 (2018).

46. Ortuño-Costela, M. del C., Cerrada, V., García-López, M. & Gallardo, M. E. The Challenge of Bringing iPSCs to the Patient. Int. J. Mol. Sci. 20, 6305 (2019).

47. Kanchan, K. et al. Genomic integrity of human induced pluripotent stem cells across nine studies in the NHLBI NextGen program. Stem Cell Res. 46, 101803 (2020).

48. Kilpinen, H. et al. Common genetic variation drives molecular heterogeneity in human iPSCs. Nature 546, 370–375 (2017).

49. Suk, H.-J., Boyden, E. S. & van Welie, I. Advances in the automation of whole-cell patch clamp technology. J. Neurosci. Methods 326, 108357 (2019).

50. Sakurai, T. The role of cell adhesion molecules in brain wiring and neuropsychiatric disorders. Mol. Cell. Neurosci. 81, 4–11 (2017).

51. Lai, H. C. & Jan, L. Y. The distribution and targeting of neuronal voltage-gated ion channels. Nat. Rev. Neurosci. 7, 548–562 (2006).

52. Gouwens, N. W. et al. Integrated Morphoelectric and Transcriptomic Classification of Cortical GABAergic Cells. Cell 183, 935-953.e19 (2020).

53. Marín, O. Interneuron dysfunction in psychiatric disorders. Nat. Rev. Neurosci. 13, 107–120 (2012).

54. Tripathy, S. J. et al. Transcriptomic correlates of neuron electrophysiological diversity. PLOS Comput. Biol. 13, e1005814 (2017).

55. Burke, K. J. & Bender, K. J. Modulation of Ion Channels in the Axon: Mechanisms and Function. Front. Cell. Neurosci. 13, (2019).

56. Chen, N. et al. Meta-analyses of RELN variants in neuropsychiatric disorders. Behav. Brain Res. 332, 110–119 (2017).

57. Grayson, D. R. & Guidotti, A. The Dynamics of DNA Methylation in Schizophrenia and Related Psychiatric Disorders. Neuropsychopharmacology 38, 138–166 (2013).

58. Guidotti, A., Grayson, D. R. & Caruncho, H. J. Epigenetic RELN Dysfunction in Schizophrenia and Related Neuropsychiatric Disorders. Front. Cell. Neurosci. 10, (2016).

59. Ishii, T. et al. In Vitro Modeling of the Bipolar Disorder and Schizophrenia Using Patient-Derived Induced Pluripotent Stem Cells with Copy Number Variations of PCDH15 and RELN. eNeuro 6, ENEURO.0403-18.2019 (2019).

60. Corces, M. R. et al. Single-cell epigenomic analyses implicate candidate causal variants at inherited risk loci for Alzheimer’s and Parkinson’s diseases. Nat. Genet. 52, 1158–1168 (2020).

61. Bulik-Sullivan, B. K. et al. LD Score regression distinguishes confounding from polygenicity in genome-wide association studies. Nat. Genet. 47, 291–295 (2015).

62. Maity, B. et al. Regulator of G Protein Signaling 6 (RGS6) Protein Ensures Coordination of Motor Movement by Modulating GABAB Receptor Signaling. J. Biol. Chem. 287, 4972–4981 (2012).

63. Sekine, K. et al. Reelin Controls Neuronal Positioning by Promoting Cell-Matrix Adhesion via Inside-Out Activation of Integrin α5β1. Neuron 76, 353–369 (2012).

64. Bosch, C. et al. Reelin Regulates the Maturation of Dendritic Spines, Synaptogenesis and Glial Ensheathment of Newborn Granule Cells. Cereb. Cortex 26, 4282–4298 (2016).

65. Kress, G. J. & Mennerick, S. Action potential initiation and propagation: upstream influences on neurotransmission. Neuroscience 158, 211–222 (2009).

66. Berg, J. et al. Human neocortical expansion involves glutamatergic neuron diversification. Nature 598, 151–158 (2021).

67. Gabaldón, T. & Koonin, E. V. Functional and evolutionary implications of gene orthology. Nat. Rev. Genet. 14, 360–366 (2013).

68. Trubetskoy, V. et al. Mapping genomic loci implicates genes and synaptic biology in schizophrenia. Nature 1–13 (2022) doi:10.1038/s41586-022-04434-5.

69. Finucane, H. K. et al. Partitioning heritability by functional annotation using genome-wide association summary statistics. Nat. Genet. 47, 1228–1235 (2015).

70. Durinck, S. et al. BioMart and Bioconductor: a powerful link between biological databases and microarray data analysis. Bioinforma. Oxf. Engl. 21, 3439–3440 (2005).

71. Gouwens, N. W. et al. Classification of electrophysiological and morphological neuron types in the mouse visual cortex. Nat. Neurosci. 22, 1182–1195 (2019).

72. Friedman, J., Hastie, T. & Tibshirani, R. Regularization Paths for Generalized Linear Models via Coordinate Descent. J. Stat. Softw. 33, 1–22 (2010).

73. Hinrichs, A. S. et al. The UCSC Genome Browser Database: update 2006. Nucleic Acids Res. 34, D590–D598 (2006).

74. Subramanian, A. et al. Gene set enrichment analysis: A knowledge-based approach for interpreting genome-wide expression profiles. Proc. Natl. Acad. Sci. 102, 15545–15550 (2005).

75. Yu, G., Wang, L.-G., Han, Y. & He, Q.-Y. clusterProfiler: an R Package for Comparing Biological Themes Among Gene Clusters. OMICS J. Integr. Biol. 16, 284–287 (2012).

76. Robinson, M. D., McCarthy, D. J. & Smyth, G. K. edgeR: a Bioconductor package for differential expression analysis of digital gene expression data. Bioinformatics 26, 139–140 (2010).

77. Csardi, G. & Nepusz, T. The igraph software package for complex network research. 9.

